# Recombinant manufacturing of multispecies biolubricants

**DOI:** 10.1101/2024.05.05.592580

**Authors:** Marshall J. Colville, Ling-Ting Huang, Samuel Schmidt, Kevin Chen, Karan Vishwanath, Jin Su, Rebecca M. Williams, Lawrence J. Bonassar, Heidi L. Reesink, Matthew J. Paszek

**Affiliations:** Robert Frederick Smith School of Chemical and Biomolecular Engineering, Cornell University, Ithaca, NY, USA; Department of Clinical Sciences, College of Veterinary Medicine, Cornell University, Ithaca, NY, USA; Department of Materials Science and Engineering, Cornell University, Ithaca, NY USA; Institute of Biotechnology, Cornell University, Ithaca, NY, USA; Nancy E. and Peter C. Meinig School of Biomedical Engineering, Cornell University, Ithaca, NY, USA; Sibley School of Mechanical and Aerospace Engineering, Cornell University, Ithaca, NY, USA

**Author notes:** These authors contributed equally (Marshall J. Colville, Ling-Ting Huang).

**Keywords:** lubricin, mucin, biomanufacturing, lubrication

## Abstract

Lubricin, a lubricating glycoprotein abundant in synovial fluid, forms a low-friction brush polymer interface in tissues exposed to sliding motion including joints, tendon sheaths, and the surface of the eye. Despite its therapeutic potential in diseases such as osteoarthritis and dry eye disease, there are few sources available. Through rational design, we developed a series of recombinant lubricin analogs that utilize the species-specific tissue-binding domains at the N- and C-termini to increase biocompatibility while replacing the central mucin domain with an engineered variant that retains the lubricating properties of native lubricin. In this study, we demonstrate the tissue binding capacity of our engineered lubricin product and its retention in the joint space of rats. Next, we present a new bioprocess chain that utilizes a human-derived cell line to produce *O*-glycosylation consistent with that of native lubricin and a purification strategy that capitalizes on the positively charged, hydrophobic N- and C-terminal domains. The bioprocess chain is demonstrated at 10 L scale in industry-standard equipment utilizing commonly available ion exchange, hydrophobic interaction and size exclusion chromatography resins. Finally, we confirmed the purity and lubricating properties of the recombinant biolubricant. The biomolecular engineering and bioprocessing strategies presented here are an effective means of lubricin production and could have broad applications to the study of mucins in general.

## INTRODUCTION

Lubricin (also known as proteoglycan 4 or PRG4) is a large mucin-like glycoprotein that was originally isolated from the synovial fluid of diarthrodial joints.^1,2^ Lubricin is a critical boundary lubricant that maintains low coefficients of friction at the surfaces of tissues, including cartilage, tendon, and eye.^3–7^ Lubricin also functions as a potent anti-adhesive and anti-fouling agent that resists cell adhesion and absorption of proteins to tissue surfaces.^8,9^ Patients with genetic mutations that preclude synthesis of functional lubricin develop early-onset joint failure, as well as other symptoms of Camptodactyly-Arthropathy-Coxa Vara-Pericarditis (CACP) syndrome.^10,11^ Prg4 -/- mice recapitulate many of the hallmark symptoms of CACP, including cartilage degeneration, synoviocyte hyperplasia, and abnormal protein deposits on the cartilage surface, highlighting the importance of lubricin in joint health across mammals.^9^

Lubricin is comprised of a highly hydrated mucin biopolymer domain flanked on each end by globular protein domains that mediate non-covalent attachment to tissues, as well as many non-biological materials.^1^ This molecular structure enables lubricin to self-assemble into a dense, brush-like structure on tissue or material surfaces.^12,13^ The strong hydration of lubricin in the brush confers its anti-adhesive and lubricating properties. In mammals, the lubricin mucin domain is zwitterionic in nature due to alternating positively charged lysine (K) and negatively charged glutamic acid (E) residues in the polypeptide repeat sequence (KEPAPTTP in humans). Peptide chains with repeating KE motifs are highly hydrated and demonstrate ultra-low fouling properties on grafted surfaces.^14–16^ Threonine (T) residues in the repeat sequence are densely grafted with O-linked N-acetylgalactosamine glycans (O-glycans), which are often capped with negatively charged sialic acids. The O-glycans strongly interact with water molecules to further contribute to the hydration layer around lubricin and the assembled brush.^17,18^ The globular end domains of lubricin bind with high affinity to extracellular matrix proteins, including collagen II, fibronectin, and cartilage oligomeric matrix protein, to mediate noncovalent grafting to tissues.^19,20^ The end domains also contain hydrophobic patches and clusters of charged residues, as well as other chemistries, that are proposed to mediate absorption to non-biological materials, including inorganic, metallic, polymeric, cationic, anionic, and hydrophobic surfaces.^13^

Lubricin has garnered attention for diverse biomedical applications due to its impressive anti-fouling and lubricating properties coupled with its innate biocompatibility. The ability of lubricin to prevent non-specific binding of proteins to surfaces is comparable or superior to polyethylene glycol (PEG), a standard for controlling fouling of implantable medical devices, biosensors, contact lenses, surgical tools, and other devices.^21,22^ Lubricin is an attractive alternative to PEG since it is non-immunogenic, whereas treating patients with PEGylated products can lead to immune responses and the synthesis of PEG-specific antibodies.^23^ The potential for recombinant production of lubricin in mammalian cells was demonstrated nearly two decades ago.^24^ Since then, there has been significant interest in the development of recombinant lubricin as an injectable or topical treatment for multiple biomedical conditions, including osteoarthritis, rheumatoid arthritis, surgical adhesions, tendon injuries, and dry eye disease.^4,7,8,25^ However, unlocking the therapeutic potential of lubricin still necessitates the development of scalable and cost-effective bioprocessing chains for its recombinant production.

Almost all bioprocess research and development for recombinant lubricin has emphasized production in Chinese hamster ovary (CHO) cells. A potential drawback of CHO cells is that they do not fully recapitulate human O-glycosylation and may generate glycans that are immunogenic in humans.^26,27^ Native human lubricin displays a mix of core 1 and core 2 O-glycan structures, whereas recombinant lubricin from CHO cells predominantly displays core 1 O-glycan structures, some of which may be sulfated.^28,29^ Core 2 O-glycans have been shown to mediate interactions with multivalent galectins to stabilize the lubricin brush on the cartilage surface, and, therefore, may contribute to the *in vivo* functionality of lubricin.^29,30^ CMP-Neu5Ac hydroxylase activity in CHO cells allows for the glycosylation of proteins with Neu5Gc sialic acids, which are not synthesized in humans and can provoke antibody responses.^31,32^ CHO cells also possess some α1,3-galactosyltransferase activity, which is absent in humans, and can generate protein products with Galα1,3-Gal residues (α-Gal) that are known to elicit adverse anaphylaxis reactions.^33^ Human host cell production systems may be a viable alternative to CHO cells for production of lubricin with core 2 O-glycans and human glycosylation patterns. Our prior work has demonstrated lab-scale production of a lubricin-like glycoprotein in HEK293-F, a suspension-adapted, human embryonic kidney cell line.^34^ However, scalable production of recombinant lubricin in controlled bioreactors has not been demonstrated nor optimized for human cell platforms, including HEK293.

Downstream bioprocess development for recombinant lubricin has focused on purification strategies that leverage the physiochemical and biochemical properties of the mucin domain. Anion exchange chromatography (AEX), which relies on the negatively charged sialic acid residues of the mucin domain for immobilization, is typically employed as the primary capture and purification step in the isolation of recombinant and animal derived lubricin.^24,34–36^ However, sialylation of recombinant products can be highly variable at pilot and manufacturing scales^37^, which raises concerns about the possibility of unexpected performance changes in AEX-based strategies for lubricin purification. Furthermore, metabolic waste products can interfere with the sialyation of O-glycan structures, creating the potential for varying sialylation patterns depending on the specific metabolic conditions of the bioreactor.^38^ Affinity chromatography using lectins that are specific to lubricin O-glycans has been applied for lab-scale purification.^35^ However, lectin affinity chromatography is unattractive for manufacturing scale production due to the high costs of the lectins and potential for supply chain disruptions owing to the natural sourcing of lectins. As an alternative to isolation strategies based on the mucin domain, the globular end domains of lubricin have unique chemical features, including dense clusters of positively charged residues and hydrophobic properties. Whether these features can be exploited in purification of lubricin must still be assessed.

Lubricin is highly conserved across mammalian species and tissues, raising the potential for therapeutic applications of recombinant lubricin in both veterinary and human medicine. For instance, keratoconjunctivitis sicca (KCS), or dry eye disease, is a prevalent ocular condition in dogs. Early clinical tests in humans indicate that lubricin may be effective at reducing the signs and symptoms of moderate dry eye.^39^ Intraarticular injection of a recombinant lubricin-like glycoprotein with a truncated mucin domain has been reported to slow the progression of cartilage degeneration in a rodent model of osteoarthritis (OA). OA is likely the most common orthopedic problem in dogs, with some reports estimating that OA affects over 20% of dogs over 1 year of age.^40^ Horses also frequently develop OA, with one-third of all 2- and 3-year-old racehorses demonstrating evidence of fetlock joint OA.^41^ Similar to humans and dogs, OA prevalence increases with age in horses, with more than 50% of horses over the age of 15 years affected by OA, and as many as 80-90% of horses over the age of 30 years.^42–44^ As OA significantly affects quality of life and is a source of wastage in the equine industry, therapeutic application of recombinant lubricin for intra-articular OA therapies has garnered attention in both equine and small animal veterinary medicine. To date, canine, equine, and other lubricin types relevant to veterinary medicine have not been recombinantly synthesized. While the mucin domain repeat sequence is highly conserved across mammals, differences in the sequences of the globular end domains raise concerns of immunological reactions for cross-species administration of recombinant lubricin.

## MATERIALS AND METHODS

### Antibodies and reagents

The following antibodies were used: mouse anti-human lubricin IgG, clone 9G3 (Cat #MABT401, Lot #3046718, 3173814, 2965145, Sigma Aldrich), The following secondary antibodies were used: goat anti-mouse IgG (H+L) secondary antibody, DyLight™ 800 4× PEG (Cat # SA5-35521, Invitrogen) and goat anti-mouse IgG (H+L) secondary antibody, HRP (Cat # 31430, Invitrogen). Doxycycline (Cat #sc-204734, Lot # G2018, Santa Cruz Biotechnology) was used for induction of gene expression. Valproic acid (VPA, Cat # P4543, Lot #, Millipore Sigma) was used as a histone deacetylase (HDAC) inhibitor where indicated. Selection of cells was with gentamycin, G418 (Cat #10131035, Thermo Fisher). PEI MAX® - transfection grade linear polyethylenimine hydrochloride (MW 40,000) (Cat # 24765-1, Polysciences) was used for transfections. Poloxamer 188, sodium bicarbonate and L-glutamine were used as cell culture supplements.

### Cell lines and culture

Suspension adapted human embryonic kidney cells (HEK293F) were acquired directly from Thermo Fisher Scientific as an authenticated product maintained according to the manufacturer’s guidelines in either FreeStyle 293 Expression medium (Cat #12-338-018, Thermo Fisher Scientific) or HyClone CDM4HEK293 media (Cat # SH30859, Cytiva). For routine culture and cell line development, cells were cultured in FreeStyle 293 Expression medium at 37 °C, 8% CO^2^, and 90% relative humidity in Corning ProCulture glass spinner flasks (Corning, NY, USA) at 120 rpm or polycarbonate erlenmeyer flask with vent cap (Cat # 431143, 431144, 431145, 431147, 431155) at 130 RPM. To obtain viable cell counts from suspension cells, a hemocytometer with trypan blue exclusion (Cat # 15250061, Life Technologies) was used. For protein production, cells were adapted into HyClone CDM4HEK293 media, supplemented with 1 g/L of poloxamer 188, 2 g/L of sodium bicarbonate and 4 mM L-glutamine, according to manufacturer’s guidelines, and cultured at 37°C, 5% CO^2^, and 90% relative humidity.

### cDNAs and expression vectors

The cDNA for a human lubricin-like glycoprotein with 59 perfect tandem repeats of the consensus lubricin repeat sequence, KEPAPTTP, was generated by custom gene synthesis (General Biosystems, Morrisville, NC) as previously reported (Shurer et al. 2019). To generate cDNAs for rHnLub, rEqLub, and rCaLub in this work, the cDNA for the 59X KEPAPTTP repeats was excised from the original plasmid with ApaI and AccI and subcloned into pBlueScript SK as an intermediate cloning vector. Double-stranded DNA fragments encoding the native human, equine, and canine PRG4 N- and C-terminal domains were synthesized through custom gene synthesis (IDT) with appropriate overlaps for Gibson assembly into the ApaI or AccI restriction sites of the pBluescript SK vector flanking the 59 KEPAPTTP repeats. In the cDNA for rHnLub, the IgK signal peptide was used in place of the native human PRG4 signal peptide. For inducible expression, the HnLub, EqLub, and CaLub cDNAs were cloned into a custom “all-in-one” piggybac expression vector that included a cassette for inducible expression of the desired glycoprotein and a bicistronic mNeonGreen marker under the control of a minimal CMV promoter with seven tetO sequences. The vector also included a second cassette for constitutive expression of the reverse tetracycline transactivator, rtTA-M2, and a bicistronic neomycin resistance marker under the control of an EF1α promoter. The rHnLub, rEqLub, and rCaLub cDNAs were subcloned from the pBlueScript SK vectors into the BamHI/NotI restriction sites of the inducible piggybac vector. cDNA sequences were validated by Sanger sequencing and next-generation whole plasmid sequencing. For constitutive expression of glycoproteins, the cDNAs for rHnLub, rEqLub, and rCaLub were amplified by polymerase chain reaction (PCR) with appropriate overlaps for Gibson assembly into the BamHI/NotI restriction sites of a piggybac vector with a constitutive CMV promoter and a neomycin resistance cassette. The complete cDNA sequences for rHnLub, rEqLub, and rCaLub, as well as the DNA sequences for the inducible and constitutive piggybac expression vectors, are included in Supplemental Table 1.

### Generation of stable cell lines

Stable cell lines were created by co-transfection of the piggybac expression plasmids for rHnLub, rEqLub, or rCaLub with a hyperactive transposase plasmid.^34^ Transfection of cells was performed using PEI as previously described.^45^ Transfected cells were selected with G418 (1 mg/mL) for 3 continues day followed by continued selection in G418 (750 μg/mL) for up to two weeks. For inducible expression cells, cells expressing high levels of the mNeonGreen expression reporter were enriched by one or two rounds of Fluorescence Activated Cell Sorting (FACS). Prior to FACS, cells were expanded to 0.5 – 1 × 10^6^ mL^-1^ and induced with doxycycline (1 μg/mL) for 24 hours. FACs was performed on a FACSAria Fusion (BD Biosciences) or FACSMelody (BD Biosciences) by gating on the top 25% of mNeonGreen expressing cells. Collected cells were expanded and prepared for cryopreservation according to the manufacturer’s guidelines (Thermo Fisher Scientific).

### Production of recombinant glycoproteins

Cryopreserved cells were thawed and transferred to a 250 mL shaker flask containing 50 mL of fresh CDM4 media to give a final viable cell density of 0.4×10^6^ mL^-1^. Cultures were grown in a shaking incubator at 37 °C, 5% CO^2^ and 85% humidity. Once viable cell density reached 2×10^6^ mL^-1^, the culture was transferred to a 2 L flask and 450 mL of media was added. The culture was returned to the incubator and monitored until the viable cell density once again reached 2×10^6^ mL^-1^, approximately 3.5 days later. Both batch and perfusion cultures were performed in a wave-mixed bioreactor (Biostat® RM, Sartorius) which maintained pH at 7.2 ± 0.1 through the addition of 1 M sodium bicarbonate or CO^2^ in the overlay gas mixture and pO^2^ at 60% ± 5% by addition of O^2^ to the overlay gas. A constant overlay of 1:1 N2:air was used, and the rocking motion was set to 22 rocks per minute at 6°. Cultures were monitored daily by withdrawing a 2 mL sample. Viability and live cell density were checked by hemocytometer using the trypan blue exclusion method. A 1 mL aliquot of the sample was centrifuged to sediment cells, then the supernatant frozen for later analysis.

Batch cultures were initiated by fitting a 20 L Flexsafe® RM optical culture bag and filling with 3.5 L of fresh media. The seed culture was connected by sterile weld and 500 mL transferred into the bioreactor. Doxycycline was added from a 10 mg/mL ethanolic stock to a final concentration of 1 μg/mL. On day 4 an additional 4.8 L of media and fresh doxycycline were added to the culture, and the culture was stopped on day 7 and immediately harvested. Clarification was performed by allowing the cells to naturally settle for two hours without rocking, followed by depth filtration (Millistak+ 0.054 m^2^ DOHC media, EMD Millipore). The filter was first flushed with 5 L ultrapure water at a constant flux of 100 L/m^2^/h, the flush water was discarded, then the sample was processed at a constant flux of 100 L/m^2^/h and collected in a sterile singleuse liquid storage bag (Flexsafe® 2D, Sartorius).

Perfusion cultures were initiated by fitting a 2 L Flexsafe® RM perfusion culture bag (Sartorius) and filling with 0.6 L fresh media. The seed culture and doxycycline were added as before to a final volume of 1 L and concentration of 1 μg/mL. Media containing 1 μg/mL doxycycline was connected to the feed line and an empty, sterile 20 L liquid storage bag (Flexsafe® 2D, Sartorius) to the harvest line. Both feed media and harvest were maintained at 4 °C throughout the process. Perfusion and harvest rates were maintained by the bioreactor control unit. Once the culture concluded the harvested media was used without further clarification.

### Purification

All chromatography operations were performed on AKTA avant (Cytiva) FPLC systems. Cation exchange development was performed on a 6.6 mm diameter column (Omnifit, Cole-Parmer) packed with POROS™ XS (ThermoFisher) to a bed height of 9.5 cm. Briefly, a desalting column (HiPrep 26/10 Desalting, Cytiva) was equilibrated with 2 C.V. of binding buffer before loading 15 mL sample and eluting with 1.5 C.V. of binding buffer. The binding buffer was 100 mM NaCl with either 50 mM phosphate, pH 6.8, or 37.5 mM phosphate, 37.5 mM acetate, pH 5.5. The sample peak was automatically collected by peak fractionation based on the UV^280^ chromatogram. The recovered sample was then loaded onto the pre-equilibrated CEX column. The column was washed with 2 C.V. of binding buffer then the sample eluted with a 20 C.V. linear gradient ending at 1.0 M NaCl and fractions were collected every 2 C.V. For large scale purification a 50 mm diameter column was packed with 388 mL of the same resin for a bed height of 20 cm. The column was equilibrated in 20 mM phosphate pH 7.2 buffer containing 100 mM NaCl before loading 8.5 L of clarified media. The column was then washed with 2 C.V. of equilibration buffer. Flowthrough and wash were collected and retained for analysis. Impurities were eluted with 2 C.V. of buffer containing 400 mM NaCl followed by rLub elution with 800 mM NaCl. Elutions were collected as individual fractions based on the corresponding peaks in the UV^280^ chromatogram.

Solubility studies were performed in 96 well clear bottom plates (Corning) by combining 50 μL of CEX product and 50 μL of ammonium sulfate solution. The plate was incubated at 21 °C for 20 minutes with shaking and the turbidity of each well measured at 360 nm on a microplate reader (FilterMax™ F5, Molecular Devices). The plates were then centrifuged at 3750 g for 20 mins and the supernatants collected from each well. Dot blots and SDS-PAGE with silver staining were performed to assess rLub concentration and sample purity.

Small scale HIC development was performed on a 6.6 mm diameter column packed with 2.75 mL of Capto™ Butyl ImpRes (8 cm bed height). For large scale purification a 26 mm diameter was packed with 55.8 mL for a bed height of 10.5 cm. Samples were prepared by slowly adding an equal volume of 2.0 M sodium sulfate to the CEX product while constantly stirring to prevent precipitation. The column was equilibrated in 20 mM phosphate buffer, pH 7.2, containing 400 mM NaCl and 1.0 M sodium sulfate. The sample was loaded onto the column and the column washed with 5 C.V. of equilibration buffer. The sample was eluted with a linear gradient ending in 20 mM phosphate pH 7.2 without additional NaCl or sodium sulfate. Because of the low absorptivity of rLub, peak collection was impractical and fractions were collected at constant 50 mL intervals.

Size exclusion chromatography development was performed on a HiPrep 26/60 Sephacryl S-200 HR prepacked column (Cytiva). For large scale purification a 50 mm diameter column was packed with 725 mL of Sephacryl S-400 HR resin for a bed height of 37 cm. For all SEC operations the column was equilibrated in 2 C.V. of 20 mM phosphate buffer, pH 7.2, with 100 mM NaCl and sample eluted in 1.5 C.V. of the same. Samples for development were 5-15 mL, and for large scale purification 35 mL. Constant volume fractions were collected in all cases due to the difficulty of detecting rLub in the UV chromatogram.

Final, sterile formulation was prepared by dead-end membrane filtration through a 0.2 μm Supor™ Kleenpack™ capsule (Pall Corp.). The filter was flushed initially flushed with 100 mL formulation buffer at 5 psi. The collected rLub from the SEC operation was transferred to a glass container fitted with an aseptic transfer aparatus and filtered into a sterile recepticle at a constant pressure of 5 psi. Finally, 100 μL samples of the filtered product were transferred to triplicate glass culture tubes containing tryptic soy broth (TSB) or fluid thioglycollate media (FTM) (Millipore).^46^ The innoculated cultures were incubated for 14 days at 21 °C (TSB) or 35 °C (FTM) to test for aerobic or anaerobic microorganisms, respectively.

### Recombinant StcE mucinase preparation

The cDNA for StcEΔ35 (Yu, Worrall, and Strynadka 2012) was synthesized by custom gene synthesis (Twist Bioscience) and inserted into the pET28b expression vector (See Supplemental Table 1). The cDNA for catalytically inactive E447D mutant was generated through mutation of StcEΔ35 using the Q5 Site-Directed Mutagenesis Kit (Cat # E0552S, New England Biolabs) with primers 5’-TCAGTCATGACGTTGGTCATAATTATG-3’ and 5’-ACTCATTCCCCAATGTGG-3’. StcEΔ35E447D was recombinantly expressed in E. coli strain NiCo21 (DE3) (Cat # C2529H, New England Biolabs). Transformed bacteria were grown in 1L of lysogeny broth medium (10 g/L tryptone (Cat #T7293, Sigma Aldrich), 5 g/L yeast extract (Cat #RC-117, G-BIOSCIENCES), and 10 g/L NaCl (Cat # BDH9286, VWR Chemicals)) in a bioreactor (BioFlo 310, New BRUNSWICK) at 37°C, agitated at 500 RPM and sparged with 3L/min air. When an OD600 of 0.6 – 0.8 was reached, the temperature was lowered to 20°C and expression was induced with 0.3 mM IPTG overnight. Cells were harvested by centrifugation at 3,000 g for 20 minutes, resuspended in lysis buffer (20 mM HEPES (Cat #JT4018, J.T. Baker), 500 mM NaCl and 10 mM imidazole (Cat #5720, Millipore), pH 7.5) with Pierce™ Protease Inhibitor Tablets, EDTA-free (Cat # A32965, Thermo Scientific), and lysed by a pressurized homogenizer (EmulsiFlex-C5, AVESTIN). The lysate was clarified by centrifugation at 10,000 g for 45 minutes and filtering through a 0.2 μm membrane filter (Cat # 17823, Sartorius). Mucinase was purified by immobilized metal affinity chromatography (IMAC) on a GE ÄKTA explorer 100 FPLC system. The lysate was loaded onto a HisTrap HP 5mL column (Cat # 17524802, Cytiva), washed with 20 column volumes of wash buffer (20 mM HEPES, 500 mM NaCl and 20 mM imidazole, pH 7.5), and eluted with a linear gradient of 20 mM to 250 mM imidazole in buffer (20 mM HEPES and 500 mM NaCl, pH 7.5). The elution fractions containing target protein were collected and buffer exchanged on a HiPrep 26/10 desalting (Cat #17508701, Cytiva) column equilibrated with storage buffer (20 mM HEPES and 150 mM NaCl, pH 7.5). The final protein was then concentrated by using Amicon Ultra 30 kDa MWCO filters (Millipore Sigma).

### ELISA

The lubricin samples were diluted in phosphate-buffered saline (PBS) with desired dilution ratio. The lubrin standards were made by two-fold serial dilution of huSynLUB58 in PBS. The catalytically inactive stcEΔ35E447D as a capture agent was diluted in PBS to 0.005 mg/mL and added to each well of Pierce™ nickel coated plates (Clear, 96-well, Cat #15442, Thermo Scientific). The plates with capture agent were covered with adhesive film and incubated at room temperature for 1 hour with shaking, followed by a rinse with phosphate-buffered saline + 0.05% Tween-20 (PBST). ELISA Ultrablock (Cat #BUF033A, Bio-Rad) blocking solution was added to each well of the plates, covered, and incubated at room temperature for 1 hour with shaking, followed by three rinses with PBST. The lubricin samples and standards were added to triplicate wells of the plates, covered, and incubated at room temperature for 1 hour with shaking, followed by three rinses with TBST. The primary antibody was 1:5,000 diluted in PBS, added to each well of the plate, covered, and incubated at room temperature for 1 hour with shaking, followed by three rinses with TBST. The mouse anti-human lubricin IgG primary antibody was 1:5,000 diluted in PBS and added to each well of the plates, covered, and incubated at room temperature for 1 hour with shaking, followed by three rinses with TBST. The goat antimouse IgG (H+L) HRP secondary antibody was 1:5,000 diluted in PBS, added to each well of the plates, covered, and incubated at room temperature for 30 minutes with shaking, followed by three rinses with TBST. 1-Step™ ultra TMB-ELISA substrate solution (Cat # 34029, Thermo Scientific) was added to each well of the plates and incubated at room temperature until desired color intensities were reached. An equal amount of stop solution (2 N sulfuric acid in water) was added to each well to stop the reaction. The absorbance at 450 nm was measured by a plate reader.

### SDS-PAGE analysis and immunoblot

Protein samples were denatured by mixing with NuPAGE™ LDS sample buffer (4X, Cat #NP0007, Invitrogen) and NuPAGE™ sample reducing agent (10X, Cat #NP0004, Invitrogen), and heated at 95oC for 10 minutes. The denatured protein samples were separated on NuPAGE™ 3–8% Tris-acetate gels (Cat #EA03785BOX, Invitrogen) according to manufacturer’s instructions and subsequently stained with Pierce™ silver stain kit (Cat # 24612, Thermo Scientific) according to manufacturer’s instructions or transferred to nitrocellulose (0.45 μm, Cat # 88018, Thermo Scientific) membranes. Membranes were blocked with 3% (w/v) bovine serum albumin (BSA) in tris-buffered saline + 0.1% Tween 20 (TBST) for 15 minutes at room temperature. The mouse anti-human lubricin IgG primary antibody were diluted 1:2000 in TBST, incubated on membranes for 1 hour at room temperature and washed three times with TBST for 5 minutes. The goat anti-mouse IgG (H+L) DyLight™ 800 4X PEG secondary antibodies were diluted 1:10000 in TBST, incubated on membranes for 1-2 hours at room temperature protected from light and washed three times with TBST for 5 minutes. Blots were imaged on a Bio-Rad, Chemidoc HP Imaging System (Bio-Rad). Image processing was performed using Fiji ImageJ software (ImageJ, U. S. National Institutes of Health). Dot blots were performed by spotting 5 μL of sample onto a dry nitrocellulose membrane. Once the membrane was completely dry the blot was processed as above.

### Fluorescent labeling of rHnLub and Dextran

Aminooxy-derivatized Sulfo-cyanine 7.5 (Cy7.5) fluorophores were covalently conjugated to sialic acids of rHnLub using the Periodate Aniline Ligation (PAL) method.^47^ Briefly, purified recombinant lubricin was diluted in PBST to a final concentration of 1 mg/mL followed by the addition of 28.5 mM sodium periodate. The reaction was incubated for 30 minutes at 4 °C with mixing then the periodate was removed on a Zeba 7 kDa molecular weight cut-off spin desalting column (Thermo Fisher Scientific) with a mobile phase of PBST. Cy7.5 was added to the recovered fraction at a final concentration of 0.625 mM along with 110 mM aniline and the reaction was incubated at room temperature overnight. Unbound dye was removed by desalting column as before.

Cy7.5-dextran was prepared through coupling of 500 kDa MW amino dextran (Thermo Fisher Scientific) with Cy7.5 NHS ester (Lumiprobe) in phosphate buffer according to manufacturer’s protocol followed by extensive dialysis in phosphate buffer using SnakeSkin 10 kDa MWCO dialysis tubing (Pierce) to remove unconjugated dye.

### Animal injections and *in vivo* imaging

All procedures involving animals were reviewed and approved by the Institutional Animal Care and Use Committee (IACUC) at Cornell University (IACUC protocol number: 2017-0084). Ten-to twelve-week-old male Sprague-Dawley rats (Harlan Sprague-Dawley, Inc.) were housed in pairs under a standard 12-hour light/dark cycle beginning at 6 am. Rats were maintained on ad libitum tap water and low fluorescence feed (Teklad Global 18% Protein Rodent Diet, Irradiated, Cat #2918, Envigo, USA) to minimize background fluorescence during imaging. Rats were induced under general anesthesia with isoflurane, and hair was clipped from the mid-abdomen to the hindlimbs, prior to and at weekly intervals following the first intra-articular injection. Rats were randomly allocated to one of three groups: Cy7.5-rHnLub, Cy7.5-free carboxylic acid, and Cy7.5-dextran. The knees were aseptically prepared, followed by trans-patellar tendon injection of 20 uL of Cy7.5 conjugates using a 27G, 0.5-inch needle and 0.5 mL tuberculin -syringe. Anesthetized rats were imaged using a fluorescent imaging chamber (IVIS Spectrum, Perkin Elmer, USA) at the following timepoints post-injection: 0-, 6- and 12-hours; 1-, 2-, 3-, 5-, 7- and 14-days; and at weekly intervals up to 56 days or until the fluorescent signal was not detectable above background fluorescence. Rats were positioned inside the imaging chamber in a supine position such that both knees were centered within the field of view. Images of both hindlimbs were obtained through a 13.2 cm^2^ window centered over the ventral midline. The subject height was set at 3 cm, and the camera was set to a constant 2 sec exposure. Images were processed in LivingImage 4.7.2 software (Perkin-Elmer, USA) using the Fluorescence Imaging Tomography (FLIT) module. Fluorescent intensity (745 nm excitation / 820 nm emission) was measured in units of Total Radiant Efficiency (TRE, [photon/s/cm^2^/steradian]/[μW/cm2]). In between imaging time points, rats were allowed to recover and allowed ad libitum cage exercise. Following the final imaging timepoint, rats were euthanized by CO2 overdose with confirmation via diaphragmatic puncture.

Images were processed in commercial software (LivingImage 4.7.2; Perkin-Elmer, USA). Each fluorescent image was combined with an overview photograph to provide anatomical context for measured fluorescence. Regions of interest (standardized 2 cm x 3 cm) were selected over the left and right knees on the combined images to limit fluorescent signal quantification to those regions while excluding the ventral abdomen. The fluorescence signal was integrated over a 2 cm by 3 cm oval region of interest (ROI) centered over the knee. Total signal intensity of the non-injected right knee was designated as background signal and subtracted from the total signal from the injected left knee. The results were fitted with a biexponential decay model using the Solver add-in for Microsoft Excel™.

### 3D Transillumination and micro-CT Imaging

The 3D transillumination feature of the IVIS Spectrum (Perkin Elmer, USA) system was used to generate a 3-dimensional reconstruction that allowed for contextualization of fluorescent signal tissue distribution following intra-articular injection. A 20-week-old female rat cadaver (n=1) was clipped from the mid-abdomen to the hindlimbs, and 20 μL of Cy7.5-rHnLub was injected intra-articularly into the left knee via a trans-patellar approach, followed by IVIS imaging as described above. Fluorescent excitation and emission filters were set at 745 nm and 820 nm respectively, the subject height was set at 3 cm, and the camera was set to auto exposure. Images were processed in LivingImage 4.7.2 software (Perkin-Elmer, USA) using the Fluorescence Imaging Tomography (FLIT) module. The resulting 3D dataset was combined with micro-CT imaging taken by the *in vivo* X-Ray micro-CT imaging system (SkyScan 1276, Bruker, USA) to provide skeletal context for the fluorescent signal by scanning the same rat cadaver at 40 μm/voxel with 100 kV and 200 μA source voltage and current, respectively. The scan was acquired using 170 ms exposures with 285 projections through a 228 degree angular spread. The optical transillumination images were combined with the CT data using the Bruker Coreg v3.0 module for alignment, DragonFly 4.1.0.647 (Object Research Systems (ORS), Inc., Montreal, Canada) for 3D visualization, and Adobe After Effects 17.5.1 and Adobe Media Encoder 14.6 (Adobe, Inc., CA, USA) for video rendering.

### O-Glycan profiling of rLub

All reagents were purchased from Sigma unless otherwise mentioned. Purified human rLub (600 μg) was denatured by heating at 100 °C for 5 min. The denatured proteins were subsequently treated with 19 mg of sodium borohydride (NaBH4) in 500 μL of 50 mM sodium hydroxide (NaOH) solution at 45 °C for 18 h (Fukuda 2001). The samples were cooled, neutralized with 10% acetic acid, passed through a Dowex H+ resin column, and lyophilized with borates removed under the stream of nitrogen. The glycans were permethylated for structural characterization by mass spectrometry using previously reported methods (Shajahan et al. 2017). Briefly, the dried eluate was dissolved with dimethyl sulfoxide (DMSO) and methylated by using methyl iodide and NaOH–DMSO base (prepared by mixing DMSO and 50% w/w NaOH solution). The reaction was quenched with water and the reaction mixture was extracted with methylene chloride and dried. The permethylated glycans were dissolved in methanol and crystallized with α-dihydroxybenzoic acid (DHBA, 20 mg/mL in 50% v/v methanol/water) matrix. Analysis of glycans present in the samples was performed in the positive ion mode by MALDI-TOF/TOF-MS using an AB SCIEX TOF/TOF 5800 (Applied Biosystem, MDS Analytical Technologies) mass spectrometer. Permethylated glycans from the samples were infused on an Orbitrap Fusion Tribrid mass spectrometer through an electrospray (ESI) probe with HCD and CID fragmentation option for further structural confirmation. The MS1 and MS2 spectra of the glycans were acquired at high resolution by a simple precursor scan, and respective ions were selected manually for further MS/MS scanning. Assignment of glycan structures were done manually and by using Glycoworkbench software, based on the fragmentation patterns and common biosynthetic pathways.

### Frictional Characterization of rEqLub

Frictional characterization of the recombinant lubricin was performed using a previously described, custom cartilage-on-glass tribometer.^48,49^ Briefly, full thickness, 6 mm diameter, cylindrical cartilage explants were harvested from the femoral condyles of neonatal bovine stifle joints (n=5). Cartilage explants were mated against a polished glass counterface and bathed in either phosphate buffered saline (PBS), bovine synovial fluid (BSF), 1 mg/mL or 0.2 mg/mL recombinant lubricin in PBS. All explants were compressed to 30% axial strain and allowed to depressurize for 1 hour. Once the samples achieved an equilibrium normal load, the counterface was slid at a range of sliding speeds between 0.1-10 mm/s using a DC motor. These compression levels and sliding speeds were chosen based on the strong correlation of the reported friction data to clinical outcomes.^50^ The coefficient of friction, *μ*, was recorded as the ratio of shear to normal force measured by a biaxial load cell. The equilibrium coefficient of friction was calculated at the end of sliding and averaged in the forward and reverse sliding directions to give a mean value for the coefficient of friction at each speed for each lubricant.

Generation of protein structures with surface electrostatic potential and surface hydrophobicity of recombinant human PRG4 N- and C-termini. The protein structures of the N-terminus (residues 1-346) and C-terminus (residues 819-1368) of recombinant human PRG4 were generated separately by ColabFold (ColabFold v1.5.2-patch: AlphaFold2 using Mmseqs2) and output as PDB files. Supplemental Table 2 contains a detailed list of the parameters used. Based on the PDB files from ColabFold, the electrostatic potential data were generated using the APBS-PDB2PQR web server (https://server.poissonboltzmann.org) with all the parameters set to default values and pH value set to 7.5 (Supplemental Table 3). The surface electrostatic potential visualizations were generated using ChimeraX (version 1.6.1). Surface hydrophobicity visualizations were generated using the molecular lipophilicity potential (MLP) model included in ChimeraX. with coloring ranging from dark cyan (most hydrophilic) to dark goldenrod (most lipophilic).

## RESULTS

### Genetic encoding of recombinant lubricins for multiple species

We sought to develop a versatile strategy for encoding lubricin-like glycoproteins that could be used for biomedical applications in human or veterinary medicine. We previously demonstrated an algorithm-based strategy for codon optimization that permits the fabrication of cDNAs encoding the repetitive mucin biopolymer domain sequences using custom gene synthesis. This approach leverages redundancy in the genetic code to find the minimally repetitive cDNA sequence that encodes a desired repetitive polypeptide chain. Codons were optimized for 59 perfect tandem repeats of the consensus mammalian lubricin repeat sequence, KEPAPTTP, and the corresponding cDNA was produced by custom gene synthesis. cDNAs for the N- and C-terminal domains of human lubricin were also synthesized, along with the non-repetitive serine (S), proline (P), and T rich domain that is positioned between the tandem repeats and C-terminal domain in human lubricin. The cDNA blocks were combined to create a complete coding sequence for an engineered human lubricin. To test the versatility of the strategy for the design of lubricins for other species, we similarly synthesized N- and C-terminal domains corresponding to native canine and equine lubricin and combined with the 59 KEPAPTTP repeats to generate cDNAs for engineered canine and equine lubricin, respectively. Schematics for the glycoprotein designs are presented in Figure 1A.

**Figure 1:**
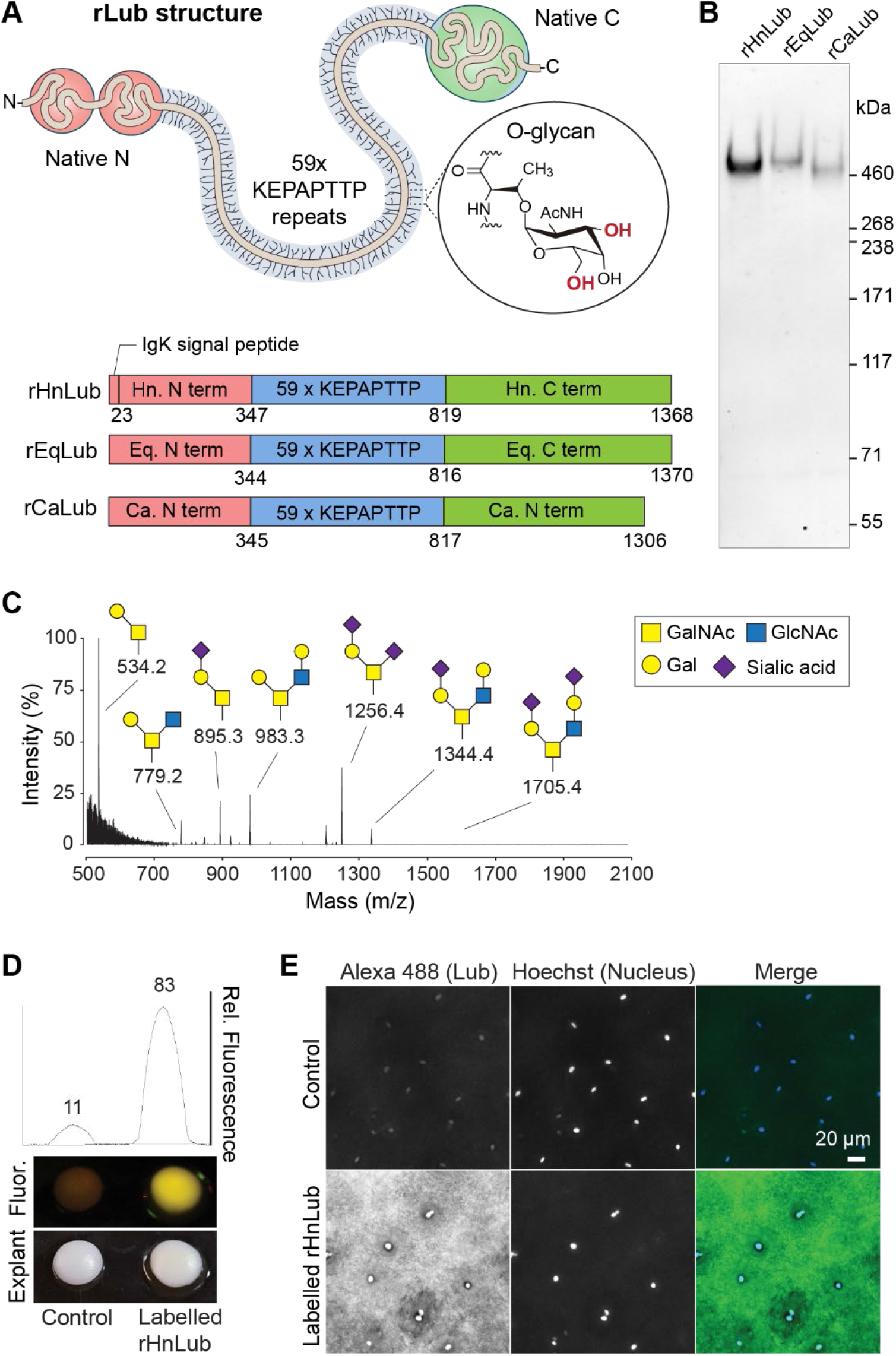
Generation of multispecies recombinant lubricins (rLub). (A) Upper: Schematic of an engineered rLub with 59 synthetic KEPAPTTP repeats flanked by the native N- and C-globular domains of native human (Hn), equine (Eq), or canine (Ca) PRG4; threonine residues in the repeats are post-translationally modified with O-GalNAc glycans that can be further extended at the 3’ and 6’ hydroxyls indicated in red. Lower: constructs for the recombinant human (rHnLub), equine (rEqLub), and canine (rCaLub) lubricin-like glycoproteins. (B) Western blot showing rHnLub, rEqLub, and rCaLub produced in HEK 293-F cells. (C) Liquid chromatography, mass spectroscopy analysis (LC-MS) of O-glycans released from rHnLub through β-elimination. (D) Photographic and fluorescent images with fluorescent quantification of bovine cartilage explants following a short incubation with either unlabeled or fluorescently labelled rHnLub. (E) Representative confocal images of the surface of the cartilage explants in D additionally labelled with Hoescht 33342 to stain the chondrocyte nuclei.

We expressed the cDNAs for the engineered human, equine, and canine lubricin glycoproteins under the control of a constitutive CMV promoter in HEK293-F cells. We recovered high molecular weight products from the media supernatants that were reactive on Western blots with the 9G3 antibody, which binds specifically to the tandem repeat sequence common in each of the lubricins (Fig. 1B). The apparent molecular weights of >400 KDa on SDS-PAGE for the multispecies lubricins were comparable to the expected molecular weights of native human, equine, and canine lubricins .^51,52^

The O-glycans of the engineered recombinant human lubricin, which we heretofore refer to as rHnLub, were profiled using liquid chromatography tandem mass spectrometry following release from the polypeptide backbone through β-elimination. We detected a mix of sialylated and non-sialylated core-1 and core-2 O-glycans, similar to the glycan profile that has been reported previously for native human lubricin isolated from synovial fluid (Fig. 1C). Although we did not conduct a glycomics analysis of the equine and canine lubricins, similar glycosylation patterns would be expected given that the tandem repeat sequences (59x KEPAPTTP) are the same for each of three glycoproteins.

### Recombinant lubricins from HEK293 have functional tissue binding activity and extended *in vivo* retention

We next tested whether the recombinant lubricin-like glycoproteins would self-assemble on the cartilage surface. To increase the production levels of rHnLub, we stably expressed it in 293-F cells with a bicistronic mNeonGreen reporter, which allowed us to isolate a high-expressing cell population using fluorescence activated cell sorting (FACS). The sialoglycans of rHnLub were metabolically labelled with azide chemical handles through supplementation of the 293-F media with N-azidoacetylmannosamine-tetraacylated (Ac4ManNAz) during production. Following purification of the rHnLub, it was fluorescently labelled through conjugation of the azido-sialoglycans with Alexa488-alkyne via copper-catalyzed click chemistry. Bovine cartilage explants were visibly fluorescent under a blue light source following a brief, 10-minute incubation with the labelled lubricin, indicating dense assembly of rHnLub on the cartilage. Confocal microscopy confirmed the assembly of the rHnLub on the surface of the cartilage. These results supported the potential for production of functional recombinant lubricin and lubricin-like products in HEK293.

Given the ability of the rHnLub to bind to cartilage surfaces, we tested whether the glycoprotein would have extended retention kinetics in joints *in vivo*, as reported for other recombinant lubricin products.^53–55^ The retention kinetics of rHnLub were evaluated in the healthy knee of Sprague-Dawley rats after a single intra-articular injection. Prior to injection, the glycans of rHnLub were fluorescently labelled using periodate oxidation to generate aldehydes on the sialic acid residues, followed by aniline-catalyzed oxime ligation with an aminooxy-derivatized sulfo-Cy7.5 (Supplemental Fig. 1). As controls, we also injected cohorts of rodents with free sulfo-Cy7.5 dye and high molecular weight (HMW) dextran (500 kDa) labelled with dye. Following intra-articular injection, signals from rHnLub, free dye, and HMW dextran were measured using IVIS imaging according to the time schedule presented in Figure 2A. Dual-mode micro-computed tomography (micro-CT) and IVIS fluorescence imaging (Figure 2B) was performed in a single cadaver to demonstrate the distribution and evaluate the depth of penetration of rHnLub after injection into the knee (Fig. 2B).

**Figure 2:**
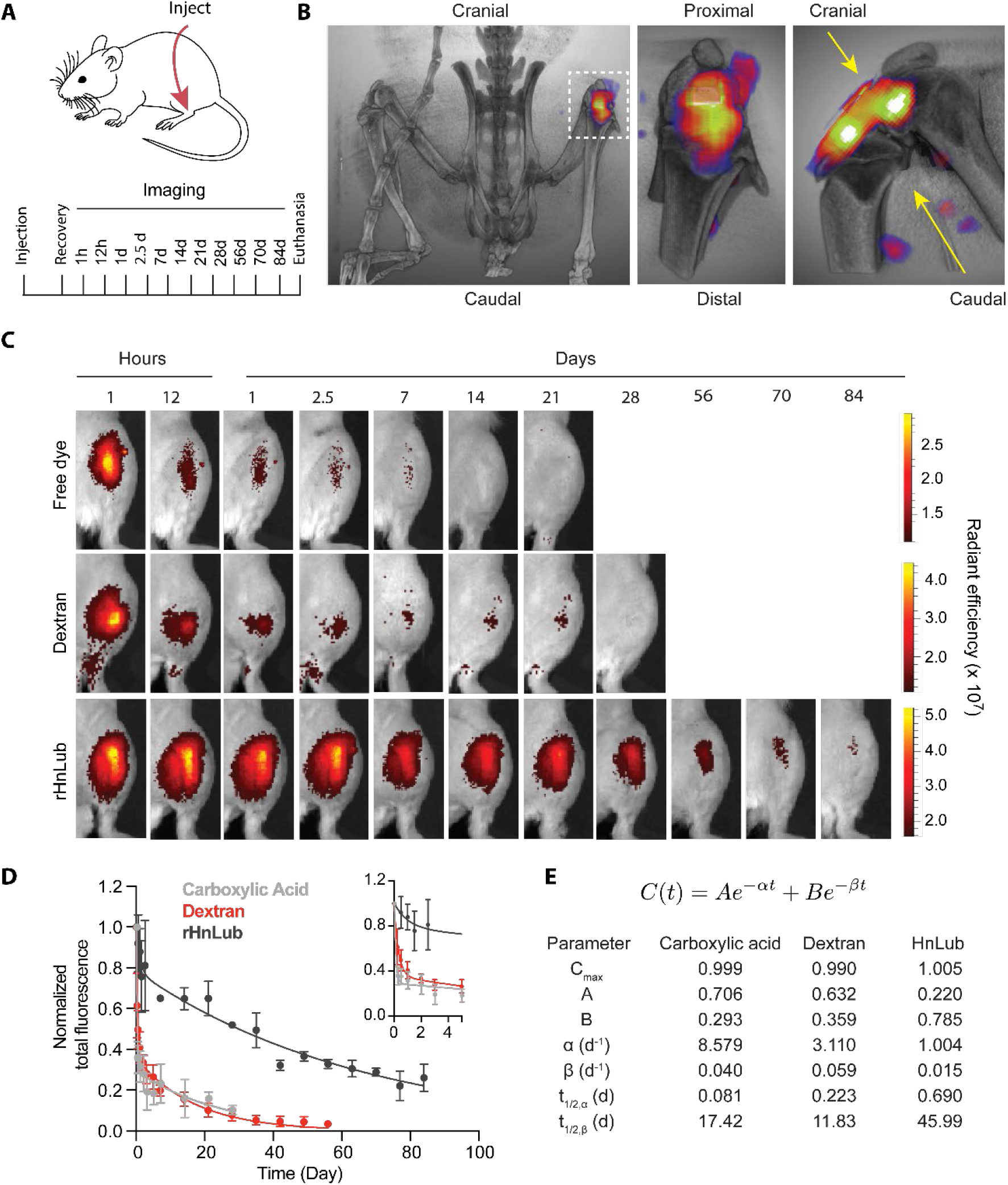
Retention kinetics of recombinant human-like lubricin following intra-articular injection. (A) Schedule for IVIS imaging of sulfo-Cy7.5 labeled rHnLub (rHnLub-Cy7.5), sulfo-Cy7.5 labelled 500 kDa dextran (Dextran-Cy7.5), and unconjugated sulfo-Cy7.5 carboxylic acid (free dye) following intra-articular injection into the knees of Sprague-Dawley rats. (B) Dual-mode micro-CT/IVIS imaging of the rat knee joint following intra-articular injection of rHnLub-Cy7.5. (C) Representative IVIS images of the rat knee at the indicated time points following intra-articular injection with free dye, dextran-Cy7.5, and rHnLub-Cy7.5. (D) Normalized total fluorescence in the rodent knee at the indicated time points following injection of free dye, dextran-Cy7.5, and rHnLub-Cy7.5. Inset shows the same curves, expanded to visualize the rapid decay in the first 5 days. (E) Model and fitted parameters from the curves in (D).

The clearances of injected rHnLub, free dye, and HMW dextran from the rodent knees were described by a standard two-compartment model (Figure 2C and 2D). Free dye and HMW dextran displayed a fast, pronounced α decay with approximately 75% of the injected compounds cleared within the first three days (Fig. 2C). The α decay was less pronounced for rHnLub, and following some minor early clearance, injected rHnLub displayed a remarkably stable β half-life of 45 days (Figure 2C and 2D). Given that metabolically labeled rHnLub bound stably to the surface of cartilage explants ex vivo, our data were consistent with the possibility that the long *in vivo* half-life of rHnLub may be attributed to its ability to bind cartilage or other joint tissues. Overall, these data suggest that recombinant intra-articular injections of rHnLub may have long-term potential therapeutic value due to the extended residence time. Previously, recombinant human lubricin and lubricin-like products with truncated tandem repeats have been reported to have extended retention kinetics following intra-articular injection in minipigs and rodents, respectively.^53–55^

### Development of a scalable production scheme for recombinant lubricins

In developing a production and purification strategy for recombinant lubricins (rLubs), our priorities were first to have a fully scalable process, second to be largely independent of glycosylation and third to be applicable across species. To this end, we chose individual unit operations in the process chain that were used in large-scale biologic manufacturing, as well as reproducible in small scale with typical laboratory equipment. Figure 3 shows a schematic diagram of our bioprocess chain, from production through packaging. The upstream operations include production in an appropriate bioreactor followed by clarification in 2 steps, sedimentation followed by filtration. The neat, clarified media was loaded onto a cation exchange chromatography (CEX) column and eluted with sodium chloride in phosphate buffer. Next, ammonium sulfate was added, and the sample was further purified in a bind-and-elute operation on a hydrophobic interaction chromatography (HIC) column. Finally, size exclusion chromatography (SEC) was used to polish the sample and accomplish a buffer exchange into the final formulation buffer before sterile filtration and packaging.

**Figure 3:**
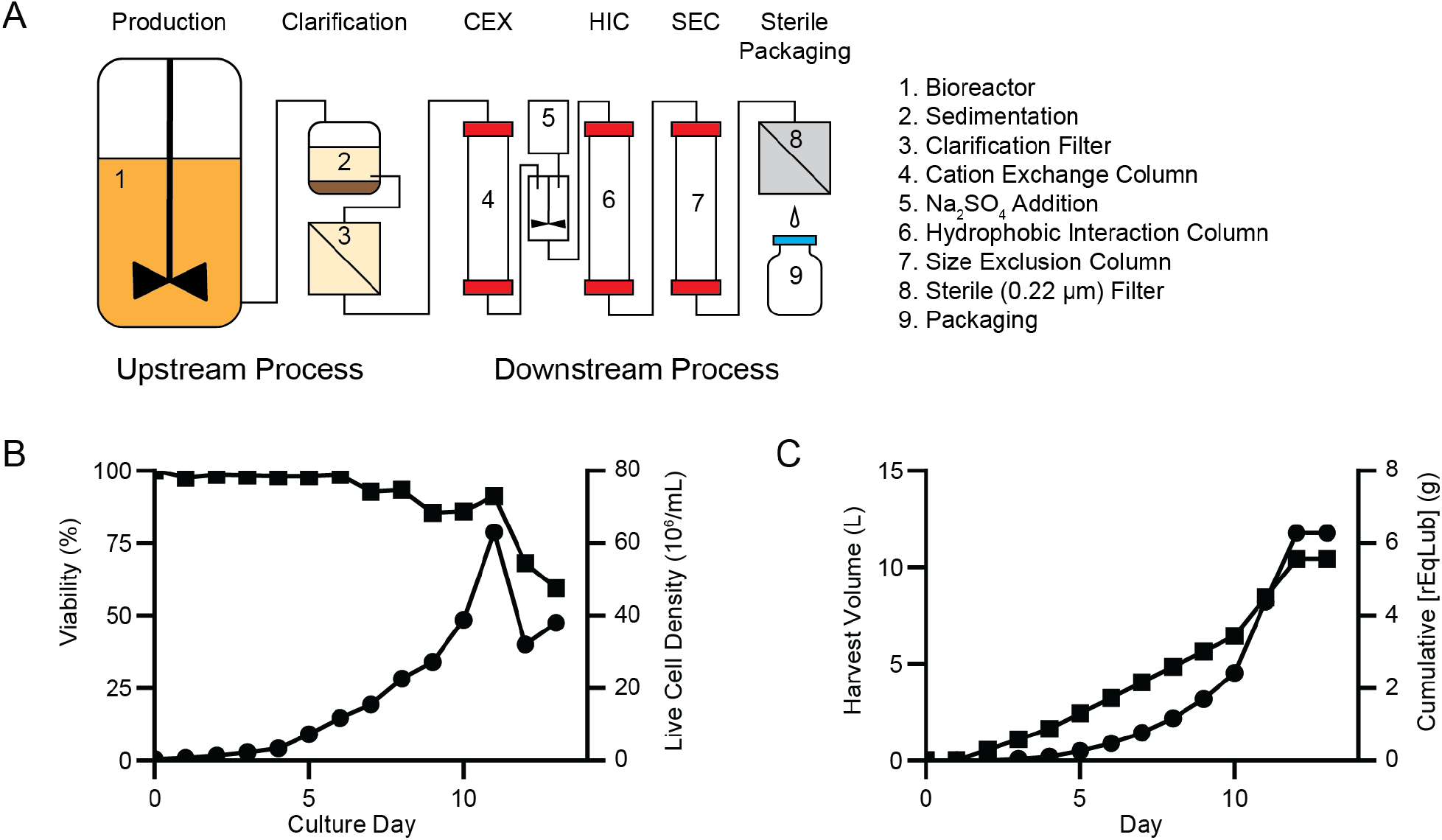
A scalable bioprocess chain to produce recombinant lubricin. (A) Schematic diagram of the complete bioprocess chain showing the minimum set of operations. (B) Cell viability (squares) and viable cell density (circles) of a 1 L rEqLub production perfusion culture. (C) Harvested volume (squares) and cumulative rEqLub production (circles) from the perfusion culture in (B).

To validate the production of rLubs with the HEK293-F production cell line in a large-scale bioreactor, we used a rocking-motion bioreactor with online pH and pO^2^ control and investigated two production schemes, batch, and perfusion cultures. Production of rEqLub was investigated as an initial test case. In an 8.9 L batch culture the viable cell density peaked at 2.73*10^6^ cells/mL on day 7 (Supplemental Fig. 2), with a final product concentration of 0.346 g/L. Pseudo-perfusion cultures in shaker flasks, in which 50% of the media was replaced every 12 hours, achieved viable cell densities of more than 10*10^6^ cells/mL. Based on these encouraging results, we chose to implement a perfusion culture scheme in the same rocking-motion bioreactor. The culture volume was maintained at 1 L for 13 days, and media was perfused at a rate of 0.55 L/day on days 2-4, 0.8 L/day on days 5-10, and 2 L/day on days 11 and 12. On day 12 the cell retention filter on the single-use culture bag became obstructed and the culture was stopped one day later. With perfusion, the viable cell density peaked on day 11 at 63*10^6^ cell/mL (Figure 3B), and the product concentration on day 12 at 0.952 g/L (Figure 3C). The rate of perfusion was increased on day 11 in response to increasing ammonia concentrations and decreasing lactate (Supplemental Fig. 3) and falling viability, however we suspect that the increased flux through the culture bag led to early fouling of the membrane. Additional optimization, including strategies, such as alternating tangential flow filtration with a replaceable membrane, could lead to higher cell densities and more efficient production.

### Product capture by cation exchange chromatography

We reasoned that rLubs could be purified based on the net positive charge of the terminal globular domains (Figure 4A, Supplemental Fig. 4). Whereas there could be a large variability in sialylation, the major source of negative charge at neutral pH and source of the low pI of fully glycosylated lubricin,^56^ the charge of the protein N- and C-terminal domains should be constant. We therefore developed a capture step to exploit the positively charged domains. Cation exchange chromatography is typically performed at pH values below the isoelectric point of the target molecule when run in a bind-and-elute mode. In small scale experiments we found that lubricin was efficiently captured at neutral pH and relatively high salt concentrations of 100 mM NaCl, and that rLub was eluted in a relatively narrow conductivity range (Figure 4B). In fact, there was very little shift in the rLub peak when the pH was increased from 5.5 to 6.8, values below and approximately equal to the pI, respectively. Furthermore, when the collected fractions were analyzed by SDS-PAGE and silver staining, we found there were fewer co-eluting impurities at pH 6.8 than 5.5 (Figure 4C,D). This data indicated that a direct capture of rLub from clarified culture media would be possible.

**Figure 4:**
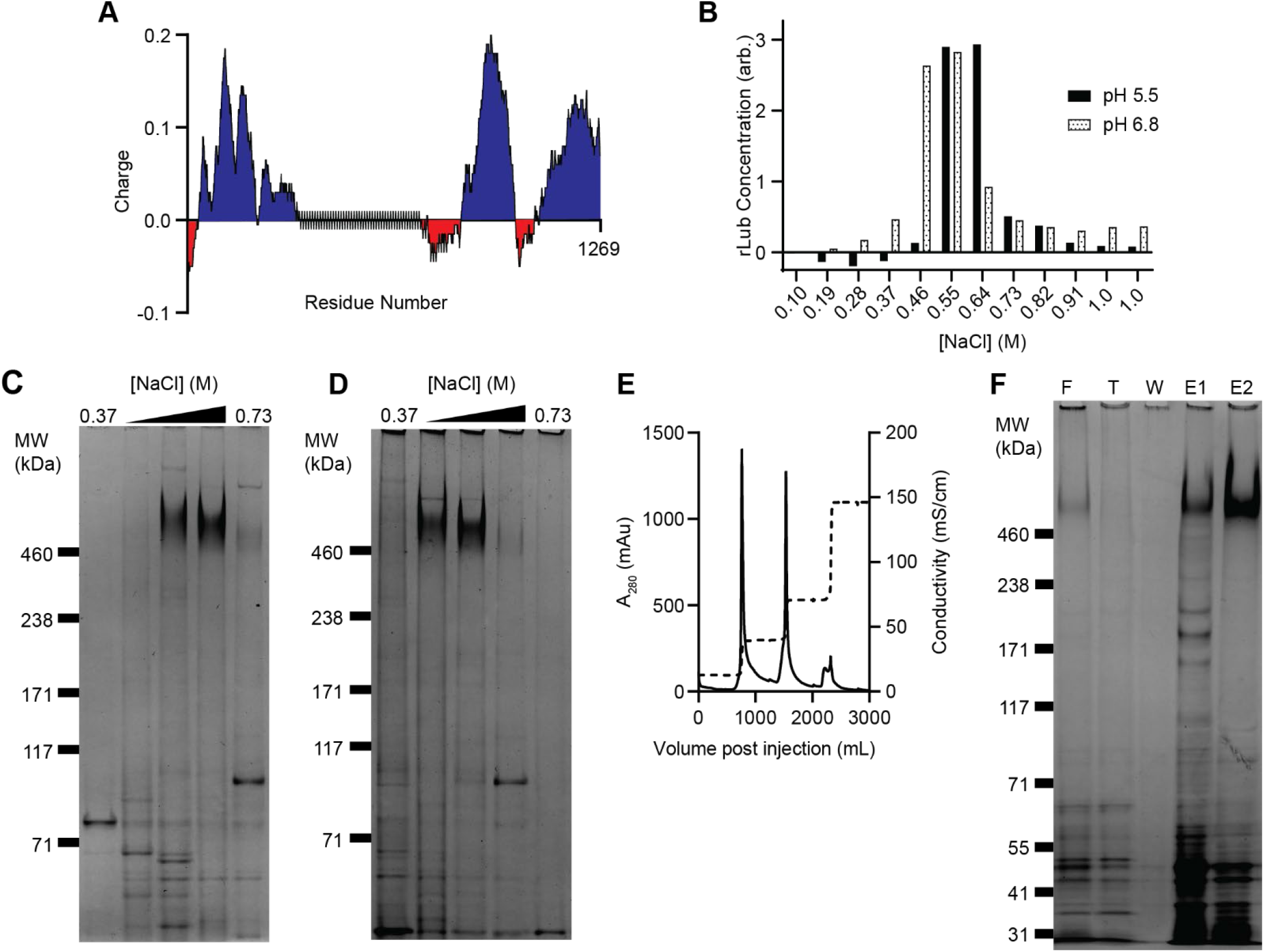
Cation exchange chromatography is a high-resolution capture operation. (A) Plot of the predicted local charge (100 AA window) along the rLub polypeptide backbone without glycosylation showing the net positive charge in the terminal, globular domains. (B) Product concentration by collected fraction during a linear gradient elution with NaCl from a cation exchange column at pH 5.5 (solid bars) or pH 6.8 (open bars). Concentration of rLub was determined by dot blot relative to the feed concentration. (C) Silver-stained gel image of the pH 5.5 separation shown in (B). The heavy bands above 460 kDa correspond to rLub. (D) Same as (C), but for the pH 6.8 separation. (E) UV absorbance (solid line) and conductivity (dashed line) chromatograms of the cation exchange step elution protocol. (F) Silver-stained gel image of the capture by step elution shown in (E). Lanes: F – feed, T – flow-through, W – column wash, E1 – 400 mM elution, E2 – 800 mM elution.

In small scale experiments, rLub was subjected to a buffer exchange by gel filtration (Sephadex G-25) into a binding buffer that matched the composition of the CEX buffer. While relatively fast, this step is prohibitively expensive as the culture volume increases. We attempted buffer exchange by tangential flow filtration (TFF), but the membrane rapidly fouled, and recovery was low. Based on the small-scale results, we reasoned that it would be possible to load rLub containing culture media directly onto a CEX column. We therefore designed a step elution protocol that would maximize recovery and purity while concentrating rLub. Figure 4E shows the chromatogram from a capture operation on a CEX column run in 20 mM phosphate buffer at pH 7.2. A column was packed with POROS™ XS strong cation exchange resin (388 mL, 20 cm bed height) and equilibrated with 100 mM NaCl in phosphate buffer before 8.5 L of clarified rLub media was loaded onto the column. The resulting load was approximately 8.3 mg rLub per mL of settled resin without detectable breakthrough. The column was then washed with 2 each of 100 mM and 400 mM NaCl to elute the bulk of the impurities. The product was recovered with a final elution in 800 mM NaCl. This protocol balances recovery with purity, as can be seen in Fig. 4F. Approximately 15% of rLub is eluted in the 400 mM fraction along with most of the impurities, leaving a concentrated product in the final elution. Further optimization may result in significant improvement in recovery (e.g., lowering the NaCl concentration in the second wash step).

### Intermediate purification by hydrophobic interaction chromatography

Although the co-eluting impurities from the capture operation are of low apparent molecular weight as shown by silver-stained protein gels (Figure 4F), we were not able to resolve them from rLub by size exclusion chromatography alone. Although lubricin forms a highly hydrated brush on the surface of tissues, the N- and C-terminal domains are predicted to display significant hydrophobic regions (Supplemental Fig. 5). We therefore investigated whether HIC could be used as an intermediate purification step. In preliminary solubility screening studies, we found that the 800 mM NaCl fraction from the capture operation was stable in up to 1.0 M ammonium sulfate (Supplemental Fig. 6) at pH values of 5 to 7. Furthermore, at neutral pH we found that most impurities began to precipitate before rLub as the concentration of ammonium sulfate was increased, indicating HIC would be an effective purification step. With this information, we developed a bind-and-elute strategy using a moderately hydrophobic HIC resin. Prior to injection, we added sodium sulfate to the sample at a final concentration of 1.0 M. The sample was then injected onto the column and eluted with a linear gradient. Figure 5A shows the UV absorbance chromatogram of the gradient elution, with the rLub peak highlighted by the grey box. The UV absorptivity of rLub is low, complicating inline detection, but the product peak was easily identified by SDS-PAGE (Figure 5B). While rLub eluted well before the major impurity peak (* in Figures 5A, B), a small, discrete population of rLub eluted with the more hydrophobic impurities. We speculate that this rLub is aggregated with the impurities or under-glycosylated, leading to its stronger interaction with the HIC resin. Given the presence of additional, rLub containing peaks in the chromatogram, additional optimization of the sodium sulfate gradient or inclusion of modifiers such as glycerol or alcohols may increase total recovery.

**Figure 5:**
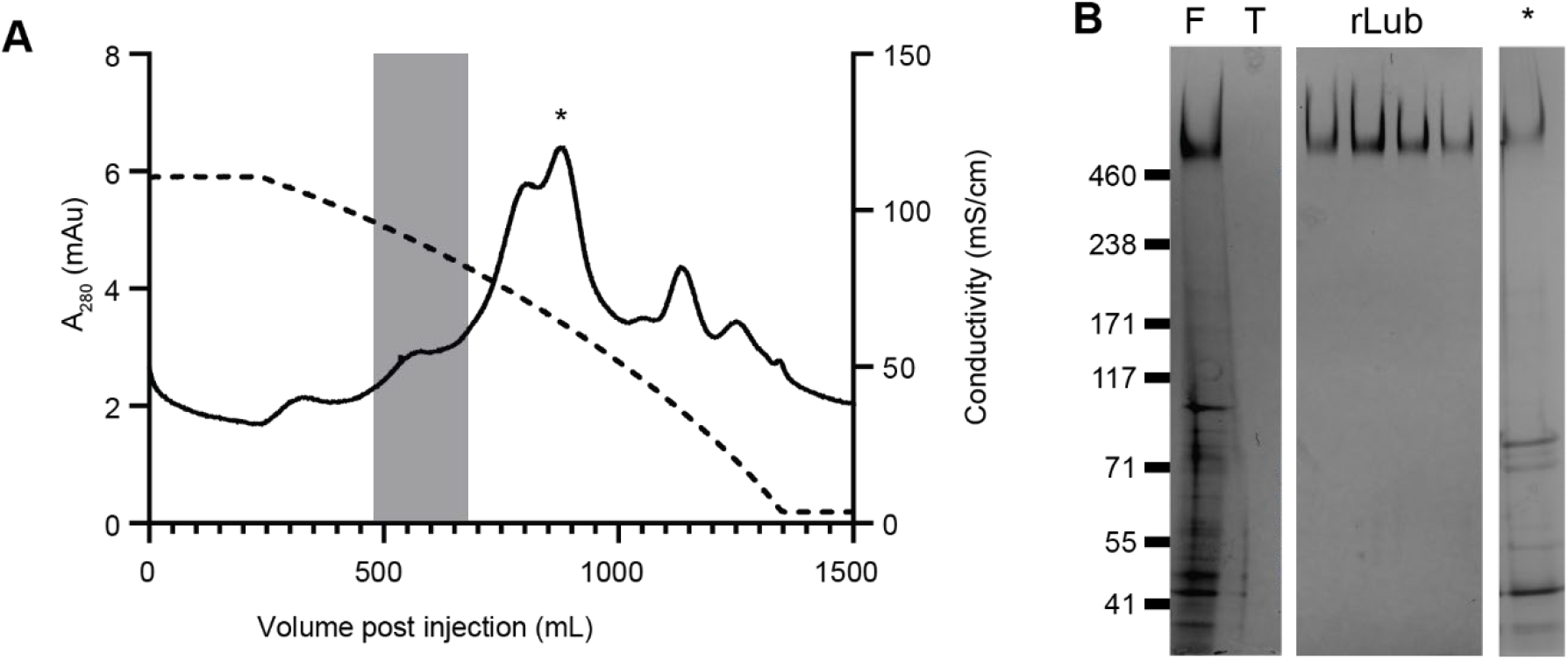
Intermediate purification with hydrophobic interaction chromatography removes most residual impurities. (A) UV absorbance (solid line) and conductivity (dashed line) chromatograms from a 1.0 M to 0 M sodium sulfate gradient elution. The peak in the shaded box corresponds to elution of the majority of rLub. The asterisk denotes the major impurity peak. (B) Silver-stained gel image of fractions from (A). Lanes: F – feed, T – flow-through. The rLub fractions are from the shaded area and asterisk from the corresponding peak in (A).

**Figure 6:**
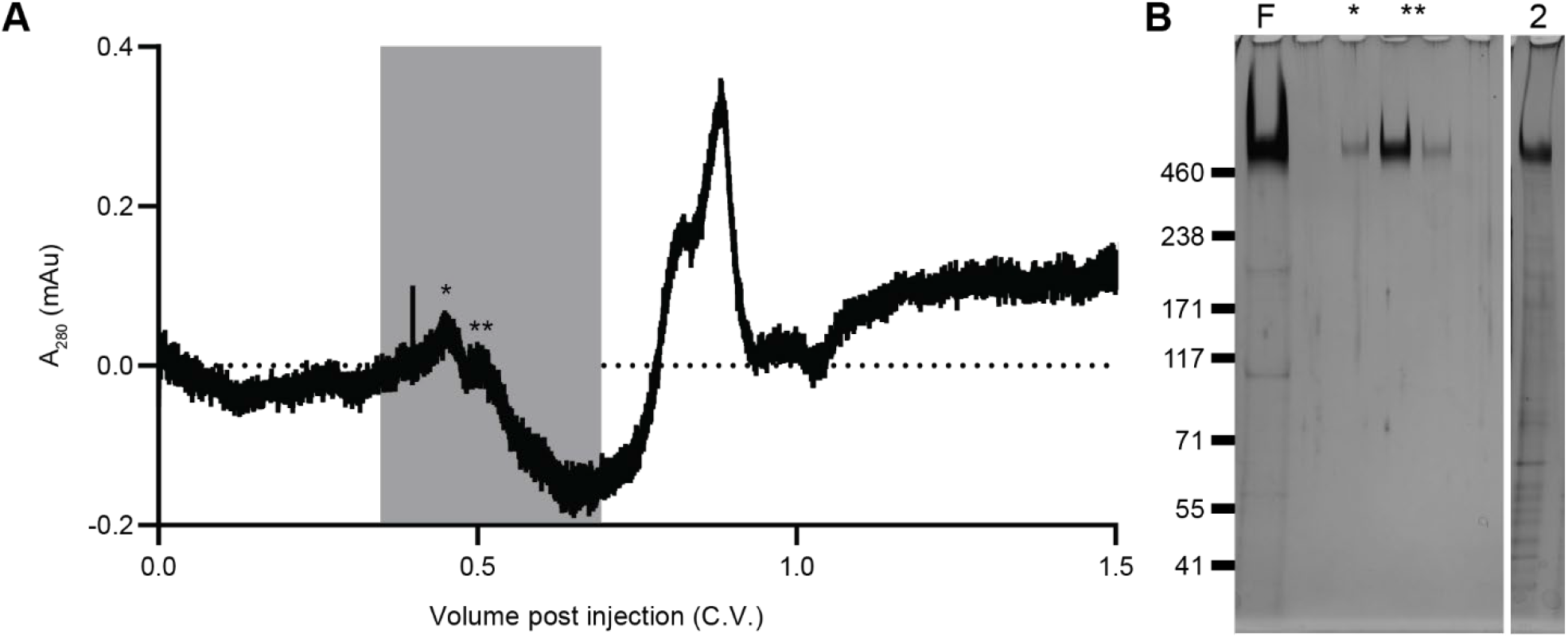
Polishing by size exclusion chromatography results in a high-purity product. (A) UV absorbance chromatogram of an SEC polishing operation on a Sephacryl S-400 column. The single and double asterisk denote peaks containing rLub. (B) Silver-stained gel image of the collected fraction from (A). Lanes: F – feed, followed by the first five fractions of the elution (shaded region in (A) with * and ** marking the lubricin containing peaks in (A)), 2 – rLub product from a 2-step process omitting intermediate purification by HIC.

### Polishing by size exclusion chromatography

The high molecular weight of rLub relative to the few remaining impurities made SEC a logical choice for a polishing operation. Figure 6A shows a typical SEC chromatogram, with the rLub containing fractions highlighted in grey. Although the rLub appears as 2 overlapping peaks in the chromatogram, they are indistinguishable by SDS-PAGE. Figure 6B shows a silver-stained protein gel of the same SEC operation, highlighting the low impurity concentration in the feed and complete absence of corresponding bands in the rLub containing fractions. The final lane, labeled “2” in Figure 6B, is from a two-step purification wherein the 800 mM NaCl fraction from the CEX operation was loaded directly onto the SEC column without the intermediate HIC step. The product of this two-step protocol retained a low concentration of impurities that made it unsuitable for *in vivo* studies, whereas we were unable to detect any residual impurities when the full protocol was followed. Furthermore, SEC not only purified the product to injection-grade, but it also allowed us to perform a buffer exchange to remove the high salt concentration in the feed material. We ran the polishing operation with phosphate buffered saline as the mobile phase, resulting in a product that was ready for sterile filtration and packaging without the need for a final diafiltration step.

### Validation of cartilage lubrication by processed recombinant lubricin

Frictional characterization of the novel recombinant rEqLub revealed it lubricated articular cartilage as effectively as native lubricin. Compared to saline, rEqLub at 1 mg/mL and BSF had a significantly lower coefficient of friction across the range of tested sliding speeds (Figure 7A, n=4-6, p < 0.001). While rEqLub at 0.2 mg/mL had a lower coefficient of friction than the saline control, it did not lubricate cartilage as effectively as the 1 mg/mL group. Like other recombinant forms of lubricin, rEqLub lubricated cartilage in a dose-dependent manner.^57^ However, the EC50 needed for lubrication could vary widely between recombinant lubricin and lubricin-like glycoprotein sources owing to differences in the protein structure, method of production, and purification. The reported coefficients of friction for rEqLub at 1 mg/mL in this study are within the same order of magnitude as previously reported friction data for other recombinant forms of lubricin.^57^ Interestingly, at the lowest sliding speed, rEqLub at 1 mg/mL and BSF have nearly identical coefficients of friction (Figure 7B, n=4-6, p = 0.09).

**Figure 7:**
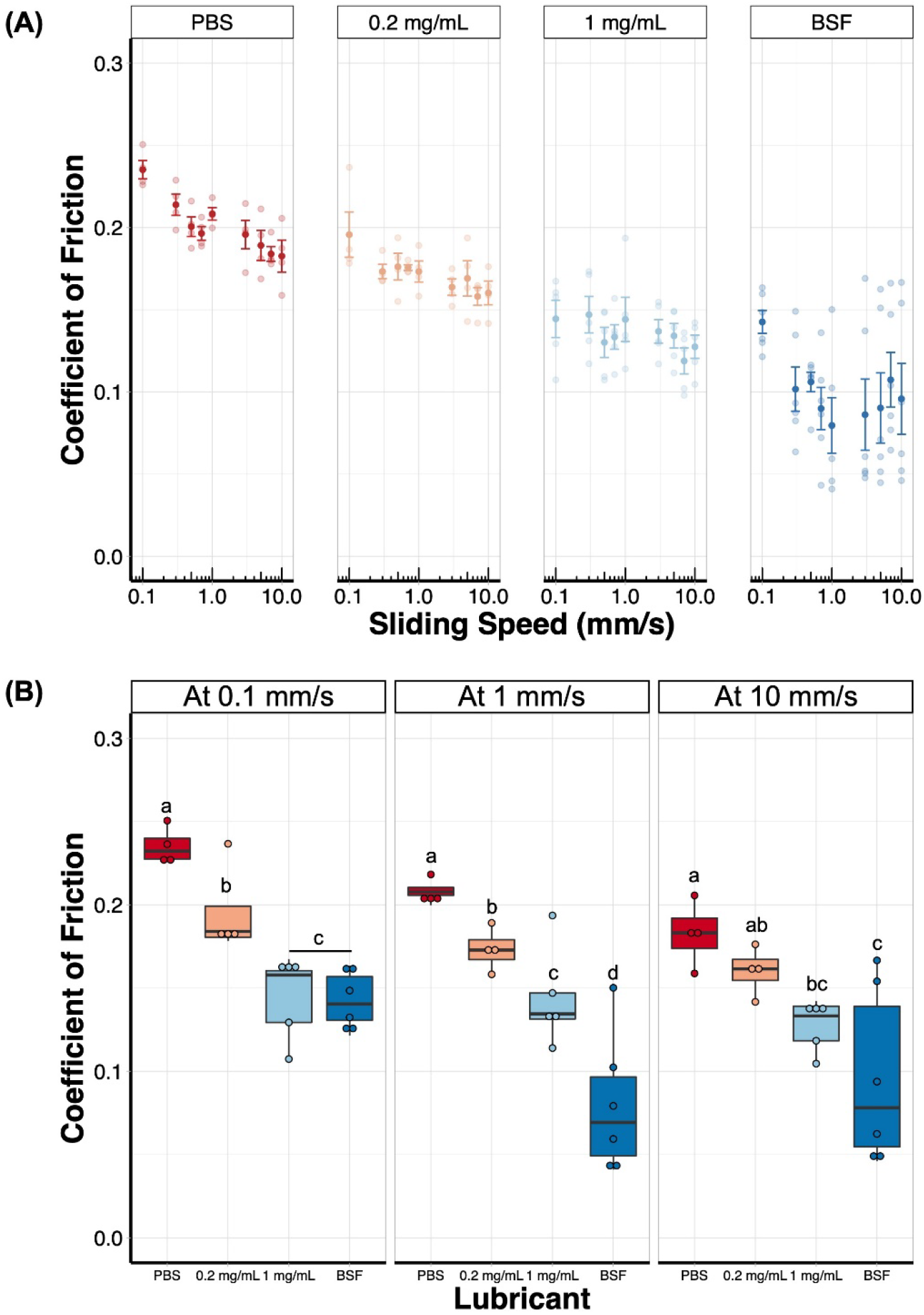
Recombinant lubricin is an effective cartilage lubricant. (A) Coefficient of friction as a function of sliding speed. Cartilage explants were maintained in a bath of either PBS, 0.2 mg/mL rEqLub, 1 mg/mL rEqLub or bovine synovial fluid (BSF) (n= 5). (B) Coefficients of friction at 0.1, 1, and 10 mm/s as a function of lubricant.

## DISCUSSION

The glycoprotein lubricin is a potent boundary lubricant that reduces sliding friction throughout the body, facilitating pain-free, gliding motion in joints, eyes, tendon sheaths and many other structures. In this study we have demonstrated an engineered, recombinant lubricin analog, designed for production in human cells, and demonstrated its potential as a long-lasting intraarticular injectable. Through the application of codon scrambling and optimization, we have designed a cDNA specifically to produce a recombinant lubricin analog that features 59 perfect repeats of the KEPAPTTP sequence, the consensus sequence for the tandem repeats (TR) in native mammalian lubricin. Furthermore, the codon optimization, scrambling, and stable integration of the cDNA into the human-derived HEK293-F production cell line demonstrated in this study indicate that the rLubs can be reproducibly manufactured across a spectrum of scales. Through replacement of the tissue-binding N- and C-terminal domains we have produced lubricin analogs with species-specificity. Presumably, this approach could be extended to other species as additional applications arise.

One of the defining characteristics of lubricin is its ability to form a highly hydrated brush on the surface of cartilage, critical to its lubricating properties. Our *in vitro* experiments using equine cartilage explants demonstrate our lubricin analog retains the tissue binding properties of native lubricin. Furthermore, *in vivo* studies wherein rat stifles were injected with a fluorescent derivative of our lubricin analog showed a clearance half-life of approximately 46 days versus 12 days for a fluorescent dextran of similar molecular weight. Although further studies are required to resolve the binding, localization, and clearance mechanics more precisely in the joint space, the extended retention of rLub and lack of adverse reaction to injection demonstrate its potential as an intra-articular injectable compound.

Given the numerous potential therapeutic applications of lubricin and lubricin-inspired biolubricants in the treatment of musculoskeletal, ophthalmic, and other biomedical applications, there is a great need for scalable, efficient production methods. A variety of methods have been reported to purify native lubricin from natural sources as well as recombinant versions from cultured cells. These methods include collection as the retentate when synovial fluid is passed through a 0.22 μm filter,^58^ heparin-affinity chromatography,^57^ anion exchange chromatography,^9,30,34^ and others. Here, we have detailed a scalable bioprocess chain that exploits the electrostatic and hydrophobic properties of the N- and C-terminal domains of lubricin which are not subject to O-glycosylation and therefore more homogeneous. Much attention has been given to the highly sialylated mucin domain and high molecular weight of lubricin with respect to purification strategies; however, the non-obvious cation exchange and hydrophobic interaction chromatography unit operations we have presented overcome any variability and polydispersity in the product. This allowed us to design an efficient 3-step purification scheme with materials and equipment already used in industrial-scale production of biologics.

Perhaps the most surprising operation, we have shown that a strong cation exchange resin is capable of capturing rLub from clarified culture media without dilution, pH adjustment, or buffer exchange despite the neutral pH and high salt content. Canonically, binding to a cation exchange column is done at pH values below the pI of the target molecule. We hypothesize that the spatially distinct nature of the positively charged globular domains from the negatively charged mucin domains makes this operation possible. Further experiments with rLub fragments could confirm this hypothesis; however, that is beyond the scope of the present work. Similarly, rLub’s structure and high degree of O-glycosylation make HIC in bind-and-elute mode an unusual choice for purification. Indeed, a flow-through operation at lower sodium sulfate concentration would be favorable for simplicity and recovery; however, we found impurities that eluted prior to rLub from the HIC column and the present approach allowed us to achieve our target purity. Finally, we were able to sterile filter rLub purified by the 3-operation bioprocess chain we present with high recovery, but recovery was compromised when the HIC operation was omitted. From this observation and the fact that the residual impurities coeluted with rLub from the SEC operation we concluded that the low recovery was due to membrane fouling by the impurities and not rLub.

Finally, we tested whether rLub was an effective boundary lubricant. Given the unprecedented nature of our purification protocol, it was possible that we were selecting for an under-glycosylated, more hydrophobic product. To validate the functionality of our product, we tested its ability to lubricate cartilage explants. Given the complexity of synovial fluid, we were surprised to find that, at low sliding speeds, a 1 mg/mL solution of rLub alone was a similarly potent lubricant to synovial fluid. Thus, we concluded that our production methods and bioprocess chain are an effective means of generating a high-quality lubricin analog suitable for use in future *in vivo* studies.

## CONCLUSIONS

Here, we have demonstrated that species specific recombinant lubricin analogs represent novel biolubricants, retaining the tissue binding and friction-reducing properties of native lubricin. We presented a new approach to the purification of recombinant lubricin that exploits the protein-specific properties of the globular domains and is less dependent on the O-linked glycosylation that is typically the basis of product capture and purification. There is a need in the field for efficient, scalable production strategies for mucins. Although the high degree of glycosylation is a defining characteristic of mucins, we have demonstrated that conscientious design of the bioprocess chain can lead to efficient approaches that do not require specialized affinity operations or expensive, low yield operations with poor scalability. Utilizing industry-standard equipment, we have validated our process in 10 L cultures, yielding gram-scale quantities as proof-of-concept. We expect that further refinement and expertise in an industrial application would increase productivity, recovery and economy of the process, further enabling the study of lubricin and lubricin-like glycoproteins in therapeutic applications and biomaterials science.

## ASSOCIATED CONTENT

### Supporting Information

Culture density and viability; nutrient and metabolite levels; rLub solubility studies; predicted surface hydrophobicity images; cDNA sequences.

## AUTHOR INFORMATION

### Author Contributions

All authors contributed to the design of experiments, interpretation of results, and preparation of the manuscript. J.S., H.L.R. and M.J.P. designed the lubricin cDNAs. S.S., L.H. and M.J.C. developed the cell lines. M.J.C and L.H. developed and tested the processes for production and purification of the recombinant products. S.S., K.C. and H.L.R. performed the injections and rodent imaging experiments, and R.M.W. performed the micro-CT imaging and 3D image reconstruction. K.V. performed the tribology experiments. M.J.C. developed the ELISA for lubricin quantification. †These authors contributed equally.

## ACKNOWLEDGMENTS

We thank V. Weaver and J. Lakins (U.C.S.F.) for the original piggybac “all-in-one” expression vector that was further modified in this study. We also thank D. Putnam for his advice and helpful discussions concerning the bioprocess design and potential pitfalls when scaling.

