## Supplemental figures and data for "Recombinant manufacturing of multispecies biolubricants"

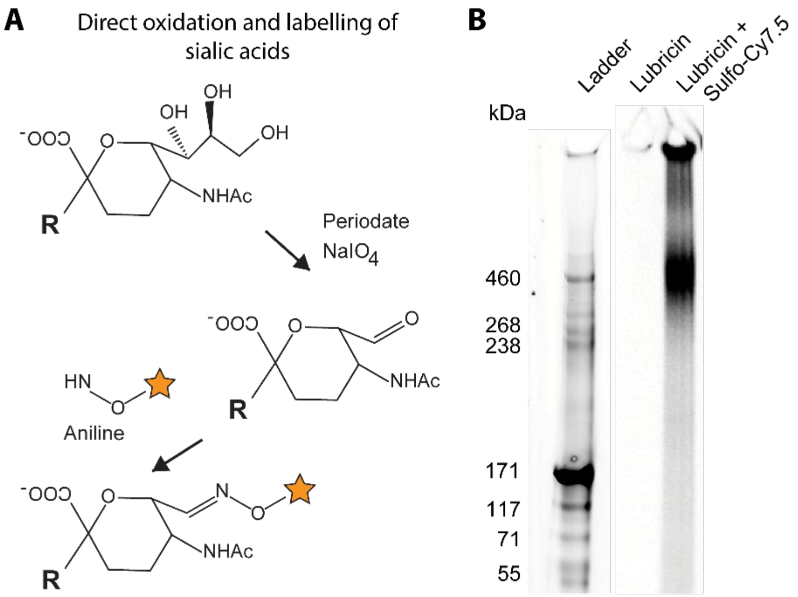


**Supplemental Figure 1: Fluorescent labeling of sialic acid containing glycans by periodate oxidation and aniline catalyzed ligation.** (A) Simplified reaction scheme for glycan-specific labeling. (B) Fluorescent blot of the Sulfo-Cy7.5 labeled rHuLub. Samples of unlabeled and labeled rHuLub were separated by SDS-PAGE on a 3-8% TRIS-acetate gel then transferred to a nitrocellulose membrane under standard conditions. The membrane was washed in TBST then imaged using a Chemidoc MP (Biorad).

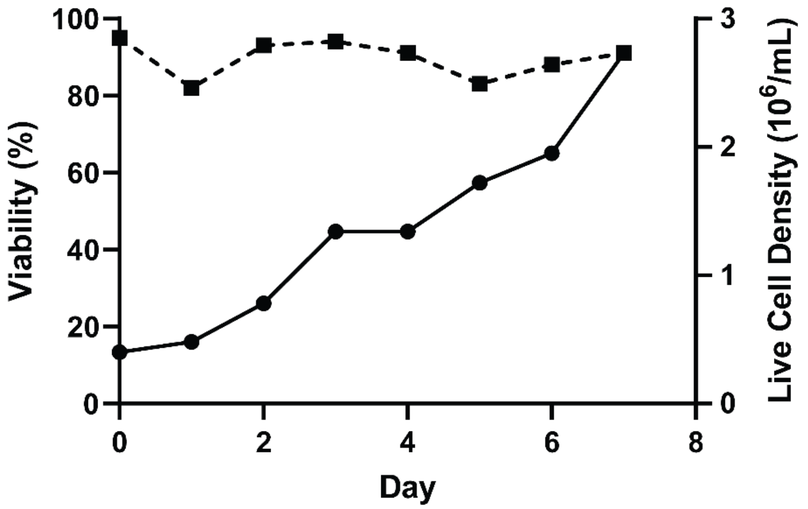


**Supplemental Figure 2: Batch production of rEqLub in a rocking motion bioreactor culture.** rEqLub producing HEK293-F cells were grown for 7 days in the presence of 1 µg/mL doxycycline. On day 4 the culture volume was increased from 4 to 8.8 L by addition of fresh media containing 1 µg/mL doxycycline. Viable cell density and viability were measured by the trypan blue exclusion method and cell counting on an improved Neubauer hemocytometer.

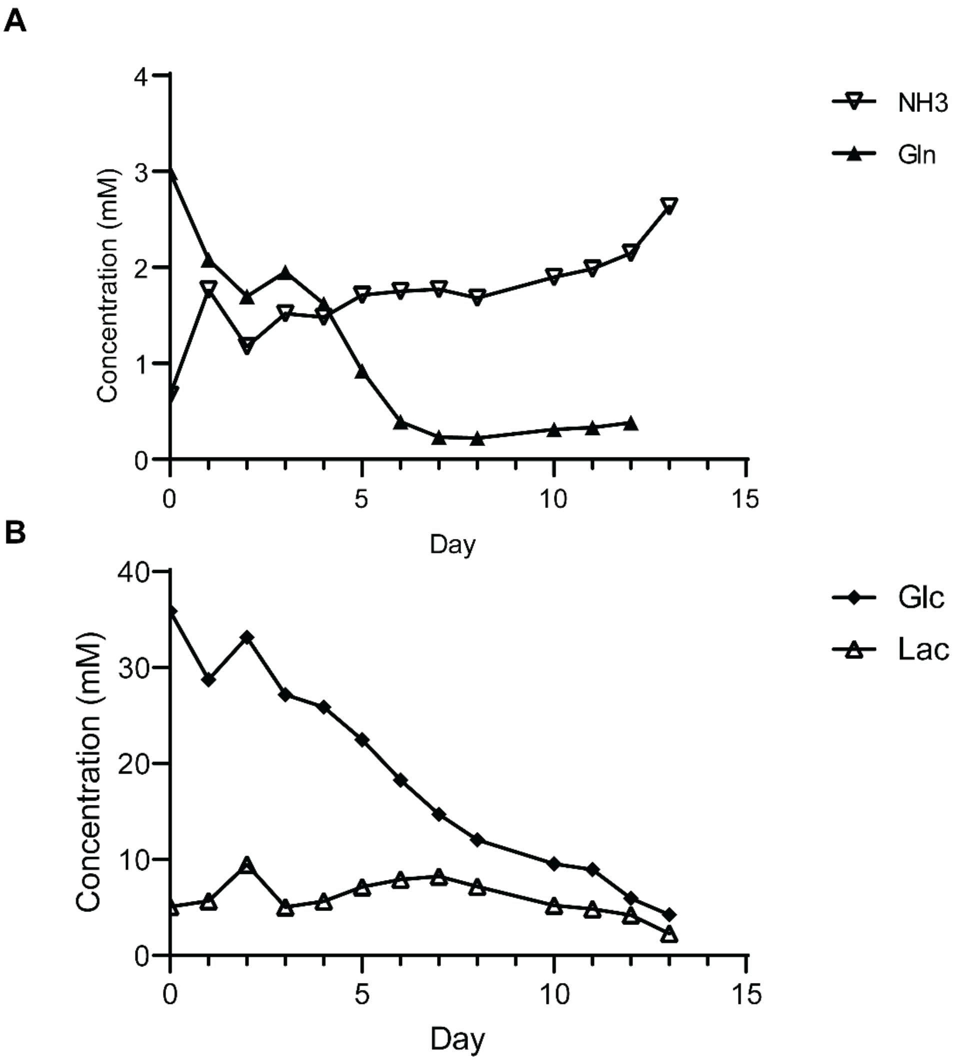


**Supplemental Figure 3: Nutrient and metabolite concentrations in perfusion culture production.** A 1 L perfusion culture was maintained by continuous addition of fresh media and harvest over a period of 13 days. Samples were taken at 24 hours intervals and analyzed for ammonia, glutamine, glucose and lactose concentrations. (A) Ammonia and glutamine concentration. (B) Glucose and lactate concentrations.

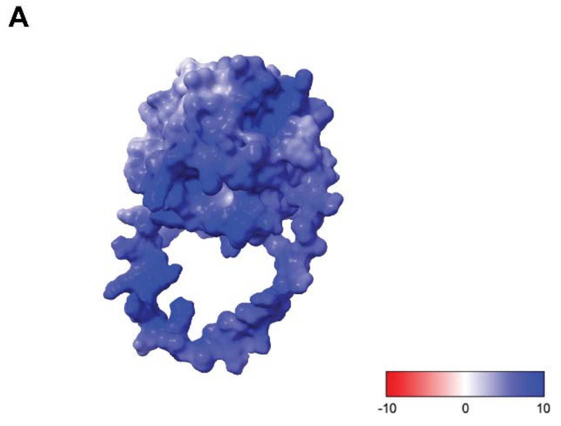


**Supplemental Figure 4: Surface electrostatic potential predictions for the tissue binding domains of rHuLub.** (A) Predicted folded structure of residues 1-84 in the C-terminal region of rHuLub. (B) Same as A for residues 1102-1368 of the N-terminal region. Electrostatic potential is displayed in units of kT/e.

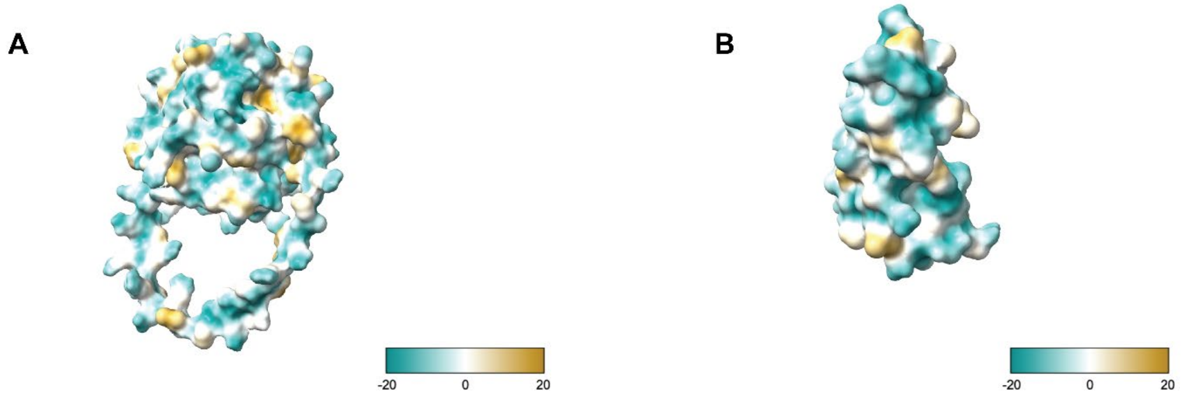


**Supplemental Figure 5: Surface hydrophobicity predictions for the tissue binding domains of rHuLub.** (A) Predicted folded structure of residues 1-84 in the C-terminal region of rHuLub. (B) Same as A for residues 1102-1368 of the N-terminal region. The molecular lipophilicity potential (MLP) is displayed on a logarithmic scale with cyan being most hydrophilic and gold most lipophilic.

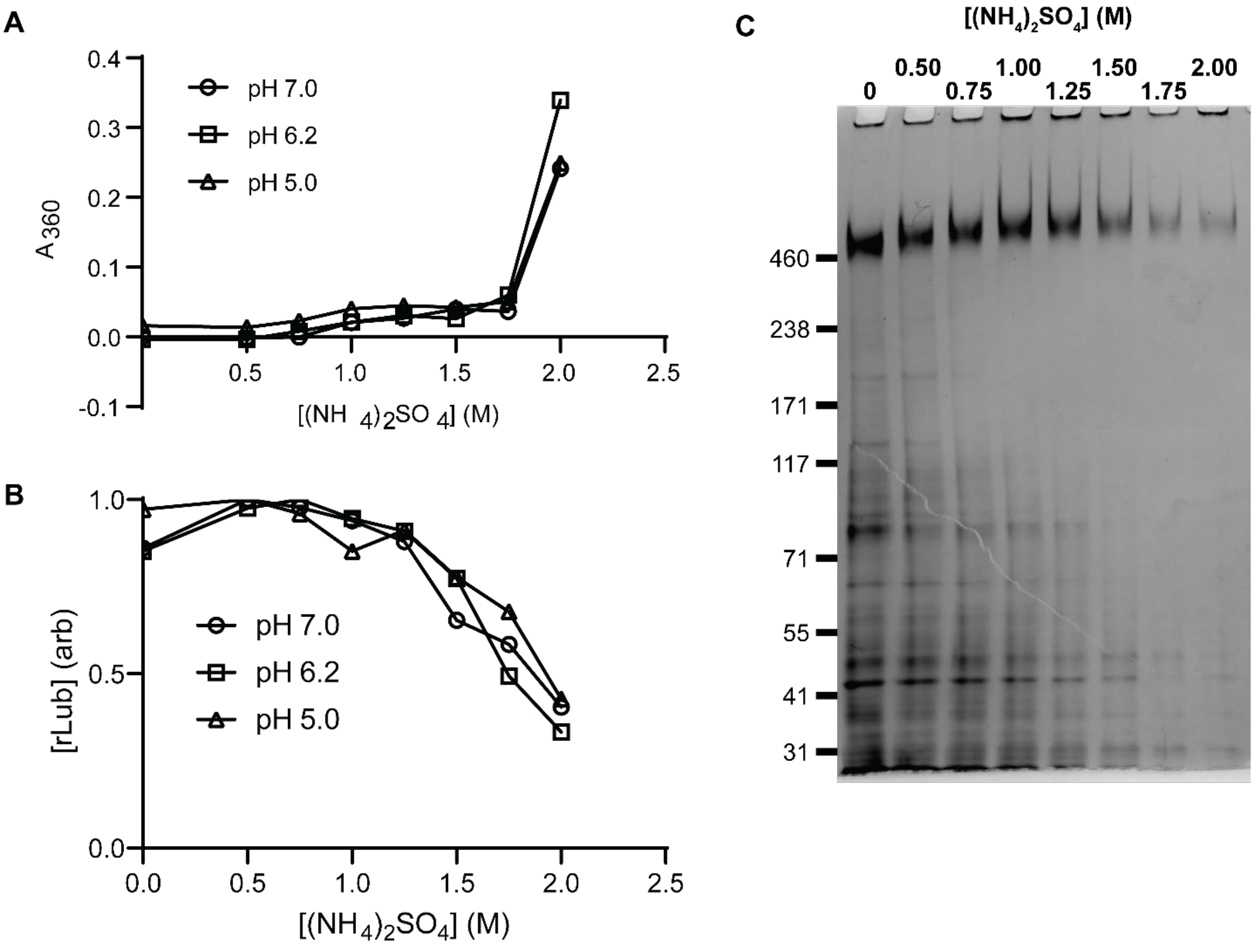


**Supplemental Figure 6: Solubility of rLub in ammonium sulfate solutions.** (A) Turbidity of recovered CEX fraction with increasing ammonium sulfate concentration determined by light scattering at 360 nm. (B) Solution concentration of rLub after addition of ammonium sulfate. Samples were centrifuged to remove precipitated components and the supernatant dotted onto a nitrocellulose membrane. rLub was detected with mouse monoclonal antibody 9G3 and DyLight 800 goat anti-mouse secondary antibody. Concentration was calculated from the fluorescence of each sample dot as measured with a ChemiDoc MP and normalized to the highest intensity.

**Supplemental Table 1. Complete genetic sequences of the species-specific rLub cDNAs and custom piggybac expression vector.**

| cDNA or Vector | Sequence |
| --- | --- |
| rHuLub | ATGGAGACAGACACACTCCTGCTATGGGTACTGCTGCTCTGGGT<br>TCCAGGTTCCACTGGTGACGGCTCCCAGGACCTGTCTAGCTGTG<br>CCGGAAGATGTGGCGAGGGCTACAGCAGAGATGCCACCTGTAA<br>CTGCGACTACAACTGCCAGCACTACATGGAATGCTGCCCCGACT<br>TCAAGAGAGTGTGCACAGCCGAGCTGAGCTGCAAGGGCAGATG<br>CTTCGAGTCCTTCGAGAGGGGCAGAGAGTGCGATTGCGACGCC<br>CAGTGCAAGAAATACGACAAGTGCTGCCCTGACTACGAGAGCT<br>TCTGTGCCGAGGTGCACAACCCCACATCTCCACCTAGCAGCAAG<br>AAGGCCCCTCCACCTTCTGGCGCCTCTCAGACAATCAAGAGCAC<br>CACCAAGCGGAGCCCCAAGCCTCCTAACAAGAAAAAGACCAAG<br>AAAGTGATCGAGAGCGAGGAAATCACCGAGGAACACAGCGTGT<br>CCGAGAATCAAGAGAGCAGCTCCAGCAGCAGCTCCTCCAGCTC<br>TAGCTCCACCATCCGGAAGATCAAGTCCAGCAAGAACAGCGCC<br>GCCAACAGAGAGCTGCAGAAAAAGCTGAAAGTGAAGGACAAC<br>AAGAAGAACCGGACCAAGAAGAAGCCCACACCTAAGCCTCCAG<br>TGGTGGATGAGGCTGGCAGCGGACTGGACAACGGCGACTTCAA<br>AGTGACCACACCTGACACCAGCACCACACAGCACAACAAGGTG<br>TCCACCTCTCCTAAGATCACCACCGCCAAGCCTATCAACCCCAG<br>ACCTAGCCTGCCTCCAAACAGCGACACCTCCAAAGAAACCAGC<br>CTGACCGTGAACAAAGAGACAACCGTCGAGACAAAAGAGACTA<br>CCACCACCAACAAGCAGACTAGTACCGACGGCAAAGAGAAAAC<br>CACCAGCGCCAAAGAGACTCAGAGCATCGAAAAGACCTCCGCC<br>AAGGATCTGGCCCCTACCTCTAAGGTGCTGGCCAAGCCAACACC<br>AAAGGCCGAGACAACCACAAAGGGCCCTGCTCTGACAACCCCT<br>AAGGAGCCAGCACCCACAACGCCGAAGGAACCAGCGCCCACGA<br>CCCCTAAAGAACCAGCTCCTACAACGCCCAAGGAACCGGCGCC<br>AACAACGCCTAAGGAACCGGCACCAACAACACCCAAAGAGCCC<br>GCCCCCACTACTCCTAAAGAACCGGCTCCAACTACACCGAAGG<br>AACCTGCCCCGACAACCCCAAAGGAACCAGCCCCTACAACCCC<br>TAAAGAGCCAGCGCCAACCACGCCCAAAGAACCTGCGCCGACT<br>ACCCCGAAAGAGCCGGCACCCACTACGCCCAAAGAGCCGGCCC<br>CCACAACCCCGAAGGAACCGGCTCCGACGACACCAAAGGAGCC<br>TGCGCCCACTACACCCAAGGAGCCTGCACCAACCACTCCCAAG<br>GAGCCAGCTCCCACAACACCAAAGGAACCCGCGCCCACCACGC<br>CAAAAGAGCCAGCACCTACAACACCTAAGGAACCTGCTCCAAC<br>CACCCCAAAGGAGCCCGCACCTACGACTCCCAAGGAACCCGCT<br>CCAACGACGCCTAAGGAGCCGGCACCTACCACTCCAAAGGAGC<br>CAGCCCCGACTACTCCGAAGGAGCCTGCCCCAACTACTCCCAAA<br>GAGCCAGCCCCCACGACTCCTAAGGAACCAGCACCAACGACAC<br>CGAAAGAACCCGCTCCCACGACGCCGAAAGAACCTGCCCCTAC |

GACACCCAAAGAACCAGCCCCAACAACTCCTAAAGAGCCGGCT
CCCACTACCCCTAAGGAGCCAGCGCCTACGACCCCAAAAGAGC
CTGCACCGACAACGCCAAAGGAACCTGCACCCACCACCCCTAA
GGAACCCGCACCAACTACCCCAAAAGAACCTGCACCTACTACT
CCAAAGGAACCGGCCCCTACCACCCCCAAGGAACCTGCGCCAA
CTACGCCGAAAGAGCCCGCGCCAACGACTCCGAAAGAACCAGC
GCCGACAACTCCAAAAGAGCCCGCTCCGACCACACCGAAAGAG
CCTGCTCCCACCACACCAAAAGAACCAGCACCGACCACTCCTA
AGGAGCCTGCTCCTACTACGCCTAAAGAACCTGCTCCGACTACA
CCTAAAGAACCCGCGCCTACCACGCCTAAAGAGCCTGCGCCTA
CAACTCCCAAAGAACCCGCACCGACTACGCCAAAAGAACCGGC
CCCAACGACCCCGAAAGAACCGGCACCGACGACTCCAAAAGAA
CCCGCCCCAACCACACCTAAAGAGCCCGCACCCACGACACCTA
AGGAGCCCGCTCCTACCACACCCAAGGAACCAGCTCCAACAAC
CCCCAAAGAGCCTGCCCCCACCACTCCGAAGGAACCCGCCCCT
ACTACACCAAAAGAGCCGGCGCCTACTACCCCCAAAGAACCGG
CGCCCACAACTCCGAAAGAGCCAGCTCCGACAACACCGAGCGA
AGTGTCTACCCCTACAACCACCAAAGAGCCAACCACCATCCAC
AAGAGCCCCGACGAGTCTACACCTGAGCTGTCTGCCGAGCCTAC
TCCTAAGGCTCTGGAAAACAGCCCCAAAGAACCCGGGGTGCCC
ACCACAAAAACACCAGCCGCCACAAAGCCCGAGATGACCACCA
CAGCCAAGGACAAGACCACCGAGCGGGACCTGAGAACAACCCC
TGAAACCACAACCGCCGCTCCAAAGATGACAAAAGAAACCGCC
ACAACCACCGAGAAAACAACCGAGAGCAAGATCACCGCCACCA
CAACACAAGTGACCTCCACCACCACTCAGGACACCACACCTTTC
AAGATCACAACCCTCAAGACCACTACACTGGCCCCAAAAGTGA
CGACCACAAAGAAAACCATCACCACGACCGAGATCATGAACAA
GCCCGAGGAAACCGCTAAGCCCAAGGACAGGGCCACCAACAGC
AAGGCCACCACACCAAAGCCACAGAAGCCTACAAAGGCCCCTA
AGAAGCCAACCAGCACAAAAAAGCCCAAGACCATGCCTAGAGT
GCGGAAGCCTAAGACAACCCCAACACCTCGGAAGATGACCAGC
ACTATGCCCGAGCTGAACCCCACCTCTAGAATCGCCGAAGCCAT
GCTGCAGACCACCACTAGACCCAATCAGACCCCTAACAGCAAG
CTGGTGGAAGTGAACCCCAAGTCCGAAGATGCCGGCGGAGCTG
AAGGCGAGACACCTCATATGCTGCTGAGGCCCCACGTGTTCATG
CCCGAAGTGACCCCTGACATGGACTACCTGCCAAGAGTGCCCA
ACCAGGGCATCATCATCAACCCTATGCTGAGCGACGAGACAAA
CATCTGCAACGGCAAGCCCGTGGACGGCCTGACCACACTGAGA
AATGGAACCCTGGTGGCTTTCCGGGGCCACTACTTTTGGATGCT
GAGCCCTTTCAGCCCTCCATCTCCTGCCAGACGGATCACAGAAG
TGTGGGGCATCCCTTCTCCAATCGACACCGTGTTCACCCGGTGC
AACTGCGAGGGCAAGACATTCTTCTTCAAGGACAGCCAGTATTG
GCGGTTCACCAACGACATCAAGGACGCCGGCTATCCCAAGCCA
ATCTTCAAAGGCTTCGGAGGCCTGACCGGCCAGATTGTGGCTGC
TCTGTCTACCGCCAAGTACAAGAACTGGCCCGAGAGCGTGTACT
TCTTTAAGAGAGGCGGCTCCATCCAGCAGTACATCTACAAGCAA

| | |
|---|---|
| | GAGCCCGTGCAGAAGTGCCCCGGAAGAAGGCCAGCTCTGAATT<br>ACCCCGTGTACGGCGAGACTACCCAAGTGCGGAGAAGAAGATT<br>CGAGAGAGCCATCGGACCCAGCCAGACACACACCATCAGAATC<br>CAGTACAGCCCCGCCAGACTGGCCTACCAGGATAAGGGCGTGC<br>TGCACAACGAAGTGAAAGTGTCCATCCTGTGGCGGGGACTGCC<br>CAATGTGGTCACAAGCGCCATCAGCCTGCCTAACATCAGAAAG<br>CCCGACGGCTACGACTACTACGCCTTTAGCAAGGACCAGTACTA<br>CAACATCGACGTGCCCAGCAGAACCGCCAGAGCCATCACAACA<br>AGATCCGGCCAGACACTGAGCAAAGTGTGGTACAACTGTCCTT<br>GA |
| rEqLub | ATGGAGTGGAAAATCCTGCCTATTTACCTTCTCCTGTTGCTGAG<br>TATATTCTCCATCCAGGAGGTTTCAAGCCAAGACCTTTCTAGTT<br>GCGCTGGTCGGTGTGGGGAGGGATACTCTCGGGATGCGACTTG<br>CAACTGCGATTTTAATTGTCAATACTACATGGAATGTTGTCCGG<br>ACTTTAAGAAAGTCTGTACATCTGAATTGTCTTGTAAAGGCCGC<br>TGTTTCGAGAGTTTCGAAAGGGGGCGAGAATGCGATTGCGATG<br>CTGACTGTAAGAAATACGGTAAGTGTTGTTCAGATTATGAAAGC<br>TTCTGCGAGGAAGTCCATAATCCTACGTCTCCGCCGAGTTCCAA<br>GACAGCTCCCCCGCCTCCAGGGGCCAGCCAGACTATCAAGAGT<br>ACAGCTAAACGGTCACCAAAGTCAAATAAGAAAAAAACTAAAA<br>AAGTTATCGAGAGTGAAGAGATCATAGAAGAACACAGTGTGTC<br>CGAGAATCAGGAGTCATCTTCCAGCTCTAGCTCAAGTTCATCTA<br>CCATCCGCAAGGTTAAGTCTAGCAAAAACTCAGCAGCGAACAG<br>AGAACTCAAAAAGAAGCCTAAGGTCAAGGATTCTAAAAAAAAA<br>CGAACCCCGAAAAAAAAACCGACGCCTGAGCCACCAGTCATAG<br>ACGAGGCCGGGAGTGGTTTGGATAACGGAGACTTCATGTTGATT<br>CCCACCCCGAAAATTCCAACCACGCAAAGAAATAAGGTGACGA<br>CATCACCAAAGATTACAACGGTAAAACCAATTAACCCCAAGCC<br>TTCCCTTCCTCCCAATTCCGACACGTCAAAAGAGACCACTAGCA<br>CACCTAATAAAGAAACTACGGTCGAGACCAAGGAGACCGAGAT<br>CACAAACAAGGAGACTTCTACAAGCGCCAATGAAAAGACTACG<br>AGCGCCAGGAAGAGTACAGAGAAAACATCCGACAAAGATTTTG<br>CTCCGGCCAGCGAAGTACCTGCAAAAAGTACCCCTAAGGCTGA<br>AACCACCACAAAGGGCCCTGCTCTGACAACCCCTAAGGAGCCA<br>GCACCCACAACGCCGAAGGAACCAGCGCCCACGACCCCTAAAG<br>AACCAGCTCCTACAACGCCCAAGGAACCGGCGCCAACAACGCC<br>TAAGGAACCGGCACCAACAACACCCAAAGAGCCCGCCCCCACT<br>ACTCCTAAAGAACCGGCTCCAACTACACCGAAGGAACCTGCCC<br>CGACAACCCCAAAGGAACCAGCCCCTACAACCCCTAAAGAGCC<br>AGCGCCAACCACGCCCAAAGAACCTGCGCCGACTACCCCGAAA<br>GAGCCGGCACCCACTACGCCCAAAGAGCCGGCCCCCACAACCC<br>CGAAGGAACCGGCTCCGACGACACCAAAGGAGCCTGCGCCCAC<br>TACACCCAAGGAGCCTGCACCAACCACTCCCAAGGAGCCAGCT<br>CCCACAACACCAAAGGAACCCGCGCCCACCACGCCAAAAGAGC<br>CAGCACCTACAACACCTAAGGAACCTGCTCCAACCACCCCAAA<br>GGAGCCCGCACCTACGACTCCCAAGGAACCCGCTCCAACGACG |

CCTAAGGAGCCGGCACCTACCACTCCAAAGGAGCCAGCCCCGA
CTACTCCGAAGGAGCCTGCCCCAACTACTCCCAAAGAGCCAGC
CCCCACGACTCCTAAGGAACCAGCACCAACGACACCGAAAGAA
CCCGCTCCCACGACGCCGAAAGAACCTGCCCCTACGACACCCA
AAGAACCAGCCCCAACAACTCCTAAAGAGCCGGCTCCCACTAC
CCCTAAGGAGCCAGCGCCTACGACCCCAAAAGAGCCTGCACCG
ACAACGCCAAAGGAACCTGCACCCACCACCCCTAAGGAACCCG
CACCAACTACCCCAAAAGAACCTGCACCTACTACTCCAAAGGA
ACCGGCCCCTACCACCCCCAAGGAACCTGCGCCAACTACGCCG
AAAGAGCCCGCGCCAACGACTCCGAAAGAACCAGCGCCGACAA
CTCCAAAAGAGCCCGCTCCGACCACACCGAAAGAGCCTGCTCC
CACCACACCAAAAGAACCAGCACCGACCACTCCTAAGGAGCCT
GCTCCTACTACGCCTAAAGAACCTGCTCCGACTACACCTAAAGA
ACCCGCGCCTACCACGCCTAAAGAGCCTGCGCCTACAACTCCCA
AAGAACCCGCACCGACTACGCCAAAAGAACCGGCCCCAACGAC
CCCGAAAGAACCGGCACCGACGACTCCAAAAGAACCCGCCCCA
ACCACACCTAAAGAGCCCGCACCCACGACACCTAAGGAGCCCG
CTCCTACCACACCCAAGGAACCAGCTCCAACAACCCCCAAAGA
GCCTGCCCCCACCACTCCGAAGGAACCCGCCCCTACTACACCAA
AAGAGCCGGCGCCTACTACCCCCAAAGAACCGGCGCCCACAAC
TCCGAAAGAGCCAGCTCCGACAACACCGAGCGAAGTGTCTACC
ACGACGACTACCATGAAACCTCCGACGACACCCAAAAATCTTG
CTGAAAGCACCCCAGAGTTCCCAGCGGAGCCAACACCCAAAGC
ACTGGAGAACTCACCCAAAGAACCGGCTGTACCGACTACGAAG
GCCCCTGAAGTAACCAAACCAGAAGTCACAACAACCGCTAAAG
ACAAGGTTACGGGAAAGGATATTCACACGATTCCCGAGATAAC
TACAGCGGCACCTAAGATAACGACCGAAACGGCCACGACAACT
GAAGAGAAAACAACGGAAAGTAAGGTGACCTCTACTATAATGC
AAGTGACCTCCACGACCGAGGATACGACGACAAGCTCCAAGAT
AACGCCTAAAGCAACGACATTGGCACCGAAAGTGATGACCGCA
ACAAAAACTACCACAACACAGGAAACGATAAACAAGCTGGAG
GAGACGACGGCTATTCCTAAGGATACGGCGACGCACAGCAAAG
TGACTACGCCAAAGCCGAAGAAGCCGACCAAAGCGCCTCGAAA
GCCGACATCCACAAAGAAACCGAAAACGCCGCGCAAGCGCAAA
CCAAAGACAACACCGATTCCCCCGAAAATCACCACCCCGACCA
CTCCTAAAAGTAACCCTACGACTTTGGCGGAAGCCATGCTTCAG
ACTACAACTTCACCTAACCAGACTCCAAATTCCGCTATGATAGA
GGTCAACCCGAAAAACGAGGACGCGGACGCTGCGGAAGGGGA
AAAGCCGCTCGTGATACTTCGACCACACGTCCTTACTCCAATCG
TCATACCGGGTCCGGACTTTCTTGTCCGCGGTCCAAACTTGGGA
ATCGGAATTAACCCCATGCTTAGCGACGAGACGAACTTGTGTAA
CGGTAAACCAGTGGACGGACTCACCACCCTGAGAAATGGAACT
CTCGTGGCTTTCAGGGGCCACTATTTCTGGATGCTCCGACCATTT
AGTCCCCCGAGTCCGCCGAGGAGAATCACCGAGGTATGGGGGA
TTCCCTCTCCTATTGATACCGTCTTCACTCGCTGCAACTGCGAGG
GAAAGACATTTTTCTTCAAGGACTCACAGTATTGGCGATTCACC

AACGACATAAAGGATGCTGGATACCCTAAATTGATTAGCAAGG
GCTTTGGGGGGCTTAGTGGCAAAATCGTGGCCGCTCTTTCAATA
GCAACGTACAAGAACAGGCCAGAGAGCGTTTATTTTTTTAAGCG
AGGGGGGCGAATACAGCAATACATCTACAAGCAAGAACCCATA
AGAAAGTGTCCAGGACGCCGACCAGCTATACATTATTCAGTTTA
CGGAGAGGCTCCTCAGATTCGGAGGAGAAGGTTCGAACGGGCC
ATAGGCCCGTCTCAGACGCACACCATCCGCATTCACTACTCCCC
CGTACGCGTATCATACCAAGACAAAGTGCCGTCCACTGACTTTC
TCCACAACGAGGTCAAAGTAAGCACCCTGTGGCGCGGACTTCC
AGACACCGTTACATCCGCCATTTCCCTTCCTAACTTGCGGAAAC
CAGACGGATACGACTATTATGCTTTTTCAAAAGACCAATATTAT
AATATTGACGTCCCGAGCCGAACTGCTCGCGCAATAACTACCCG
AAGTGGCCAGACATTGAGTAAGGTCTGGTATAACTGTCCCTAG

rCaLub

ATGCAATGGAAGATTCTCCCCATATACTTGTTGCTGCTCAGTGT
ATTCCTCATCCAACAAGTAAGTAGTCAAGATCTCCCTTCTTGTG
CAGGCAGGTGTGGAGAAGGCTATAGTCGGGATGCGATTTGTAA
TTGTGATTATAACTGCCAACATTACATGGAGTGCTGTCCGGACT
TTAAGAAAGCATGTACGGTCGAGCTCAGTTGTAAAGGGCGCTG
TTTCGAATCTTTCGCTAGAGGCCGAGAATGTGACTGCGACAGTG
ACTGCAAAAAGTACGGAAAGTGTTGCCCAGATTACGAGGACTT
TTGCGGGAGAGTACACAACCCTACTTCACCACCTTCTTCCAAAA
CTGCACCACCTTCCCCGGGGGCCTCTCAGACAATTAAGTCAACG
GCCAAACGCTCACCCAAGGCTCCGAACAAAAAAAAGACTAAGA
AGGTAATAGAGAGTGAGGAAATCACCGAGGAGCACTCTGTGTC
AGAAAACCAAGAAAGTTCTTCATCATCAAGCTCTTCTTCATCCA
CTATTCGCAAAATAAAGTCATCTAAGAACTCTGCGGCGAATAA
AGAGCTTAAAAAGAAGCCAAAAGTAAAGGATAATAAAAAGGA
GCGAACACCGAAGAAAAAGCCACCACCTGAACCCCCCGTAGTT
GATGAGGCGGGGTCAGGCTTGGACAATGGAGACATTAAATTGA
CACCCACGCCTGACATTCCTACGACTCAACGAAATAAGGTTACT
ACAAGTCCCAAATTCACCACAGGTAAGCCCATCAACCCAAAAC
CTAGTCTCCCACCGAACACCGATACGTCAAAGGAGACGTCATCC
ACTCCCAACAAGGAAACAACTGTCAAAAGTAAAGAGACACTTG
CTAACAAGGAAACCAGCAGTAAAGCGAAGGAGAAAATTACGTC
TGCTAAAGAGACTCGGTCTGCGGAGAAGACCCCAGCGAAGGAC
TTTGTGCCTACGACGAAAGCCCCTGTCAAATCTACTCCGAAGGC
GGAAAGCACTACTAAGGGCCCTGCTCTGACAACCCCTAAGGAG
CCAGCACCCACAACGCCGAAGGAACCAGCGCCCACGACCCCTA
AAGAACCAGCTCCTACAACGCCCAAGGAACCGGCGCCAACAAC
GCCTAAGGAACCGGCACCAACAACACCCAAAGAGCCCGCCCCC
ACTACTCCTAAAGAACCGGCTCCAACTACACCGAAGGAACCTG
CCCCGACAACCCCAAAGGAACCAGCCCCTACAACCCCTAAAGA
GCCAGCGCCAACCACGCCCAAAGAACCTGCGCCGACTACCCCG
AAAGAGCCGGCACCCACTACGCCCAAAGAGCCGGCCCCCACAA
CCCCGAAGGAACCGGCTCCGACGACACCAAAGGAGCCTGCGCC
CACTACACCCAAGGAGCCTGCACCAACCACTCCCAAGGAGCCA

GCTCCCACAACACCAAAGGAACCCGCGCCCACCACGCCAAAAG
AGCCAGCACCTACAACACCTAAGGAACCTGCTCCAACCACCCC
AAAGGAGCCCGCACCTACGACTCCCAAGGAACCCGCTCCAACG
ACGCCTAAGGAGCCGGCACCTACCACTCCAAAGGAGCCAGCCC
CGACTACTCCGAAGGAGCCTGCCCCAACTACTCCCAAAGAGCC
AGCCCCCACGACTCCTAAGGAACCAGCACCAACGACACCGAAA
GAACCCGCTCCCACGACGCCGAAAGAACCTGCCCCTACGACAC
CCAAAGAACCAGCCCCAACAACTCCTAAAGAGCCGGCTCCCAC
TACCCCTAAGGAGCCAGCGCCTACGACCCCAAAAGAGCCTGCA
CCGACAACGCCAAAGGAACCTGCACCCACCACCCCTAAGGAAC
CCGCACCAACTACCCCAAAAGAACCTGCACCTACTACTCCAAA
GGAACCGGCCCCTACCACCCCCAAGGAACCTGCGCCAACTACG
CCGAAAGAGCCCGCGCCAACGACTCCGAAAGAACCAGCGCCGA
CAACTCCAAAAGAGCCCGCTCCGACCACACCGAAAGAGCCTGC
TCCCACCACACCAAAAGAACCAGCACCGACCACTCCTAAGGAG
CCTGCTCCTACTACGCCTAAAGAACCTGCTCCGACTACACCTAA
AGAACCCGCGCCTACCACGCCTAAAGAGCCTGCGCCTACAACT
CCCAAAGAACCCGCACCGACTACGCCAAAAGAACCGGCCCCAA
CGACCCCGAAAGAACCGGCACCGACGACTCCAAAAGAACCCGC
CCCAACCACACCTAAAGAGCCCGCACCCACGACACCTAAGGAG
CCCGCTCCTACCACACCCAAGGAACCAGCTCCAACAACCCCCA
AAGAGCCTGCCCCCACCACTCCGAAGGAACCCGCCCCTACTAC
ACCAAAAGAGCCGGCGCCTACTACCCCCAAAGAACCGGCGCCC
ACAACTCCGAAAGAGCCAGCTCCGACAACACCGAGCGAAGTGA
CAACGACGGCTAAAGATAAAACGACCGAGAAAGACATAATTCC
AGAGATTACCACTGCTGTTCCCAAGATCACAACTCAAGAAACTG
CTACGCCAACCGAGGAGACGACTACGGAATCTAAGACCTCAAC
TACGACCCAAGTCACTTCTACTACTAGTAGCAAAAACACTCCAA
AAGCCACGACCCTCGCGCCCAAGGTGATGACAGCAACACAAAA
AACCACGACTACTGAAGAGACCATGAACAAGCCCGAAGAGACG
ACGGCAGTGCCTAAAGATACTGCAACATCAACGAAGGTAAGCA
CCCCGCGACCCCGAAAGCCAACCAAAGCACCAAAGAAACCCGC
AAGTACAAAGAAACCCAACACGATCCCTAAACGAAAAAAACCA
AAAACTACACCTACCCCGCCAAAGATGACTACGAGCACTATGC
CTAAACTCCATCCTACCTCCTCCGTTGAGGCAATGCTGCAAACT
ACAACGTCCCCCAATCAACGACCTAATTCTGAGATAGTAGAGGT
CAACCCCAACGAGGATACGGACGCGGCTGGAAAGAAACCCCAT
ATGTTCCCGCGACCTCCTGTTTTGACCCCCATATTTATCCCTGGA
ACCGACATTCTTGTGCGGGGGTCCAATCAAGATATTGCCATAAA
TCCCATGCTTTCCGACGAGACAAATCTCTGTAATGGAAAACCTG
TCGACGGATTGACAACCCTCCGAAATGGTACTATGGTGGCGTTC
CGCGGCCATTATTTCTGGATGTTGAGTCCTTCCAAACCCCCGAG
TCCTCCCCGGAAGATTACAGAGGTTTGGGGCATCCCCTCTCCCA
TAGATACCGTTTTTACGCGATGCAATTGTGAGGGTAAAACATTC
TTCTTCAAGGGCAGTCAGTACTGGCGATTCACTAACGACATCAA
GGACGCAGGCTACCCCAAACAGATCGTCAAGGGTTTCGGAGGC

TTGAATGGTCGAATTGTCGCTGCCCTGTCTATAGCTAAGTACAA
GGACCGGCCAGAGTCTGTCTATTTTTCAAGCGCGGCGGCTCAG
TGCAACAATATACTTACAAGCAAGAGCCGATAAAAAATGTAC
AGGGCGCCGGCCGGCGATTAACTACCCTGTATATGGTGAGACT
ACACAAGTGAGGCGGAGACGCTTTGAGAGGGCGATAGGCCCTT
CTCAGACGCATACCATCCGGATACACTACTCCCCTATTCGGGTT
AGCTACCAGGACAAGGGTTTCTTGCACAATGAAGTAAAAATGT
CCAGTCAATGGAGAGGTTTCCCGAACGTTGTTACCTCAGCAATT
GCGCTGCCTAACATCAGGAAGCCTGATGGTTACGACTATTACGC
GTTTTCTCGCAATCAATATTATAACATTGATGTTCCCTCCCGCAC
TGCCAGAGTTGTGACTACAAGATTTGGACGAACCCTCTCCAATA
TATGGTACAATTGCCCCTAG

Piggybac “all-in-one” vector with mNeonGreen reporter of expression

TATTCTGGGGGGTGGGGTGGGGCAGGACAGCAAGGGGGAGGAT
TGGGAAGACAATAGCAGGCATGCTGGGGATGCGGTGGGCTCTA
TGGCATCGATGCTAGTTAAAAGTTTTGTTACTTTATAGAAGAAA
TTTTGAGTTTTTGTTTTTTTTTAATAAATAAATAAACATAAATAA
ATTGTTTGTTGAATTTATTATTAGTATGTAAGTGTAAATATAATA
AAACTTAATATCTATTCAAATTAATAAATAAACCTCGATATACA
GACCGATAAAACACATGCGTCAATTTTACGCATGATTATCTTTA
ACGTACGTCACAATATGATTATCTTTCTAGGGTTAATCTAGTAT
ACGCGTGAGCAAAAGGCCAGCAAAAGGCCAGGAACCGTAAAA
AGGCCGCGTTGCTGGCGTTTTTCCATAGGCTCCGCCCCCCTGAC
GAGCATCACAAAAATCGACGCTCAAGTCAGAGGTGGCGAAACC
CGACAGGACTATAAAGATACCAGGCGTTTCCCCCTGGAAGCTCC
CTCGTGCGCTCTCCTGTTCCGACCCTGCCGCTTACCGGATACCTG
TCCGCCTTTCTCCCTTCGGGAAGCGTGGCGCTTTCTCATAGCTCA
CGCTGTAGGTATCTCAGTTCGGTGTAGGTCGTTCGCTCCAAGCT
GGGCTGTGTGCACGAACCCCCCGTTCAGCCCGACCGCTGCGCCT
TATCCGGTAACTATCGTCTTGAGTCCAACCCGGTAAGACACGAC
TTATCGCCACTGGCAGCAGCCACTGGTAACAGGATTAGCAGAG
CGAGGTATGTAGGCGGTGCTACAGAGTTCTTGAAGTGGTGGCCT
AACTACGGCTACACTAGAAGGACAGTATTTGGTATCTGCGCTCT
GCTGAAGCCAGTTACCTTCGGAAAAAGAGTTGGTAGCTCTTGAT
CCGGCAAACAAACCACCGCTGGTAGCGGTGGTTTTTTTGTTTGC
AAGCAGCAGATTACGCGCAGAAAAAAAGGATCTCAAGAAGATC
CTTTGATCTTTTCTACGGGGTCTGACGCTCAGTGGAACGAAAAC
TCACGTTAAGGGATTTTGGTCATGAGATTATCAAAAAGGATCTT
CACCTAGATCCTTTTAAATTAAAAATGAAGTTTTAAATCAATCT
AAAGTATATATGAGTAAACTTGGTCTGACAGTTACCAATGCTTA
ATCAGTGAGGCACCTATCTCAGCGATCTGTCTATTTCGTTCATCC
ATAGTTGCCTGACTCCCCGTCGTGTAGATAACTACGATACGGGA
GGGCTTACCATCTGGCCCCAGTGCTGCAATGATACCGCGAGACC
CACGCTCACCGGCTCCAGATTTATCAGCAATAAACCAGCCAGCC
GGAAGGGCCGAGCGCAGAAGTGGTCCTGCAACTTTATCCGCCT
CCATCCAGTCTATTAATTGTTGCCGGGAAGCTAGAGTAAGTAGT
TCGCCAGTTAATAGTTTGCGCAACGTTGTTGCCATTGCTACAGG

CATCGTGGTGTCACGCTCGTCGTTTGGTATGGCTTCATTCAGCTC
CGGTTCCCAACGATCAAGGCGAGTTACATGATCCCCCATGTTGT
GCAAAAAAGCGGTTAGCTCCTTCGGTCCTCCGATCGTTGTCAGA
AGTAAGTTGGCCGCAGTGTTATCACTCATGGTTATGGCAGCACT
GCATAATTCTCTTACTGTCATGCCATCCGTAAGATGCTTTTCTGT
GACTGGTGAGTACTCAACCAAGTCATTCTGAGAATAGTGTATGC
GGCGACCGAGTTGCTCTTGCCCGGCGTCAATACGGGATAATACC
GCGCCACATAGCAGAACTTTAAAAGTGCTCATCATTGGAAAAC
GTTCTTCGGGGCGAAAACTCTCAAGGATCTTACCGCTGTTGAGA
TCCAGTTCGATGTAACCCACTCGTGCACCCAACTGATCTTCAGC
ATCTTTTACTTTCACCAGCGTTTCTGGGTGAGCAAAAACAGGAA
GGCAAAATGCCGCAAAAAAGGGAATAAGGGCGACACGGAAAT
GTTGAATACTCATACTCTTCCTTTTTCAATATTATTGAAGCATTT
ATCAGGGTTATTGTCTCATGAGCGGATACATATTTGAATGTATT
TAGAAAAATAAACAAATAGGGGTTCCGCGCACATTTCCCCGAA
AAGTGCCACCGCTAGATTAACCCTAGAAAGATAGTCTGCGTAA
AATTGACGCATGCATTCTTGAAATATTGCTCTCTCTTTCTAAATA
GCGCGAATCCGTCGCTGTGCATTTAGGACATCTCAGTCGCCGCT
TGGAGCTCCCGTGAGGCGTGCTTGTCAATGCGGTAAGTGTCACT
GATTTTGAACTATAACGACCGCGTGAGTCAAAATGACGCATGAT
TATCTTTTACGTGACTTTTAAGATTTAACTCATACGATAATTATA
TTGTTATTTCATGTTCTACTTACGTGATAACTTATTATATATATA
TTTTCTTGTTATAGATATCGGGTCAGCTTGGCCTAGCGCTGTCGA
GTTTACCACTCCCTATCAGTGATAGAGAAAAGTGAAAGTCGAGT
TTACCACTCCCTATCAGTGATAGAGAAAAGTGAAAGTCGAGTTT
ACCACTCCCTATCAGTGATAGAGAAAAGTGAAAGTCGAGTTTA
CCACTCCCTATCAGTGATAGAGAAAAGTGAAAGTCGAGTTTACC
ACTCCCTATCAGTGATAGAGAAAAGTGAAAGTCGAGTTTACCA
CTCCCTATCAGTGATAGAGAAAAGTGAAAGTCGAGTTTACCACT
CCCTATCAGTGATAGAGAAAAGTGAAAGTCGAGCTCGGTACCC
GGGTCGAGTAGGCGTGTACGGTGGGAGGCCTATATAAGCAGAG
CTCGTTTAGTGAACCGTCAGATCGCCTGGAGACGCCATCCACGC
TGTTTTGACCTCCATAGAAGACACCGGGACCGATCCAGCCTCCG
CGGAATTTGATTAGCTTTATTGCGGTAGTTTATCACAGTTAAATT
GCTAACGCAGTCAGTGCTTCTGACACAACAGTCTCGAACTTAAG
CTGCAGTGACTCTCTTAAGGTAGCCTTGCAGAAGTTGGTCGTGA
GGCACTGGGCAGGTAAGTATCAAGGTTACAAGACAGGTTTAAG
GAGACCAATAGAAACTGGGCTTGTCGAGACAGAGAAGACTCTT
GCGTTTCTGATAGGCACCTATTGGTCTTACTGACATCCACTTTGC
CTTTCTCTCCACAGGTGTCCACTCCCAGTTCAATTACAGCTCTTA
AGGCTAGAGGATCCACTAGTCCAGTGTGGTGGAATTCTGCAGAT
ATCCAGCACAGTGGCGGCCGCGCCCCTCTCCCTCCCCCCCCCCT
AACGTTACTGGCCGAAGCCGCTTGGAATAAGGCCGGTGTGCGTT
TGTCTATATGTTATTTTCCACCATATTGCCGTCTTTTGGCAATGT
GAGGGCCCGGAAACCTGGCCCTGTCTTCTTGACGAGCATTCCTA
GGGGTCTTTCCCCTCTCGCCAAAGGAATGCAAGGTCTGTTGAAT

GTCGTGAAGGAAGCAGTTCCTCTGGAAGCTTCTTGAAGACAAA
CAACGTCTGTAGCGACCCTTTGCAGGCAGCGGAACCCCCCACCT
GGCGACAGGTGCCTCTGCGGCCAAAAGCCACGTGTATAAGATA
CACCTGCAAAGGCGGCACAACCCCAGTGCCACGTTGTGAGTTG
GATAGTTGTGGAAAGAGTCAAATGGCTCTCCTCAAGCGTATTCA
ACAAGGGGCTGAAGGATGCCCAGAAGGTACCCCATTGTATGGG
ATCTGATCTGGGGCCTCGGTGCACATGCTTTACATGTGTTTAGT
CGAGGTTAAAAAAACGTCTAGGCCCCCCGAACCACGGGGACGT
GGTTTTCCTTTGAAAAACACGATGATAATATGGCCACAACCATG
GCGTCCGGAATGGAGAGCGACGAGAGCGGCCTGGTTTCTAAGG
GAGAGGAGGATAATATGGCGAGTTTACCTGCAACTCACGAATT
GCATATTTTTGGATCTATAAACGGCGTTGATTTTGACATGGTTG
GCCAAGGAACAGGTAACCCGAATGACGGGTACGAAGAGTTGAA
CTTAAAGAGCACAAAGGGAGACCTTCAATTTTCGCCCTGGATAC
TGGTACCTCACATCGGGTACGGATTTCACCAATACCTGCCATAT
CCAGACGGCATGTCACCCTTCCAGGCCGCAATGGTCGATGGGTC
AGGTTACCAAGTCCACCGTACTATGCAGTTTGAAGACGGCGCGT
CCTTGACTGTGAATTACCGCTACACATACGAAGGAAGTCACATA
AAGGGCGAGGCGCAAGTGAAGGGAACAGGGTTTCCCGCAGATG
GACCTGTTATGACTAACAGCTTGACAGCAGCAGACTGGTGCCGT
TCAAAAAAGACTTACCCGAACGACAAAACTATTATTTCTACCTT
TAAGTGGTCTTATACTACAGGTAATGGGAAACGGTATAGATCA
ACTGCTAGAACCACATATACTTTCGCAAAACCGATGGCTGCAAA
CTATTTAAAGAACCAACCTATGTACGTTTTCAGAAAAACGGAGT
TAAAACATTCCAAAACAGAGCTTAACTTCAAGGAGTGGCAAAA
GGCATTCACTGATGTAATGGGTATGGATGAACTGTACAAATAGT
CTAGAGGGCCCGTTTAAACCCGCTGATCAATCGAAATGACCGA
CCAAGCGACGCCCAACCTGCCATCACGAGATTTCGATTCCACCG
CCGCCTTCTATGAAAGGTTGGGCTTCGGAATCGTTTTCCGGGAC
GCCGGCTGGATGATCCTCCAGCGCGGGGATCTCATGCTGGAGTT
CTTCGCCCACCCCAACTTGTTTATTGCAGCTTATAATGGTTACAA
ATAAAGCAATAGCATCACAAATTTCACAAATAAAGCATTTTTTT
CACTGCATTCTAGTTGTGGTTTGTCCAAACTCATCAATGCTCGA
CGATAAGCTTTGCAAAGATGGATAAAGTTTTAAACAGAGAGGA
ATCTTTGCAGCTAATGGACCTTCTAGGTCTTGAAAGGAGTGGGA
ATTGGCTCCGGTGCCCGTCAGTGGGCAGAGCGCACATCGCCCAC
AGTCCCCGAGAAGTTGGGGGGAGGGGTCGGCAATTGATCCGGT
GCCTAGAGAAGGTGGCGCGGGGTAAACTGGGAAAGTGATGTCG
TGTACTGGCTCCGCCTTTTTCCCGAGGGTGGGGGAGAACCGTAT
ATAAGTGCAGTAGTCGCCGTGAACGTTCTTTTTCGCAACGGGTT
TGCCGCCAGAACACAGGTAAGTGCCGTGTGTGGTTCCCGCGGG
CCTGGCCTCTTTACGGGTTATGGCCCTTGCGTGCCTTGAATTACT
TCCACTGGCTGCAGTACGTGATTCTTGATCCCGAGCTTCGGGTT
GGAAGTGGGTGGGAGAGTTCGAGGCCTTGCGCTTAAGGAGCCC
CTTCGCCTCGTGCTTGAGTTGAGGCCTGGCCTGGGCGCTGGGGC
CGCCGCGTGCGAATCTGGTGGCACCTTCGCGCCTGTCTCGCTGC

TTTCGATAAGTCTCTAGCCATTTAAAATTTTTGATGACCTGCTGC
GACGCTTTTTTTCTGGCAAGATAGTCTTGTAAATGCGGGCCAAG
ATCTGCACACTGGTATTTCGGTTTTTGGGGCCGCGGGCGGCGAC
GGGGCCCGTGCGTCCCAGCGCACATGTTCGGCGAGGCGGGGCC
TGCGAGCGCGGCCACCGAGAATCGGACGGGGGTAGTCTCAAGC
TGGCCGGCCTGCTCTGGTGCCTGGCCTCGCGCCGCCGTGTATCG
CCCCGCCCTGGGCGGCAAGGCTGGCCCGGTCGGCACCAGTTGC
GTGAGCGGAAAGATGGCCGCTTCCCGGCCCTGCTGCAGGGAGC
TCAAAATGGAGGACGCGGCGCTCGGGAGAGCGGGCGGGTGAGT
CACCCACACAAAGGAAAAGGGCCTTTCCGTCCTCAGCCGTCGCT
TCATGTGACTCCACGGAGTACCGGGCGCCGTCCAGGCACCTCGA
TTAGTTCTCGAGCTTTTGGAGTACGTCGTCTTTAGGTTGGGGGG
AGGGGTTTTATGCGATGGAGTTTCCCCACACTGAGTGGGTGGAG
ACTGAAGTTAGGCCAGCTTGGCACTTGATGTAATTCTCCTTGGA
ATTTGCCCTTTTTGAGTTTGGATCTTGGTTCATTCTCAAGCCTCA
GACAGTGGTTCAAAGTTTTTTTCTTCCATTTCAGGTGTCGTGAGG
AATTTCGACATTTAAATTTAATTAATCTCGACGGTATCGGTTGG
GATCTGCCACCATGTCCAGACTGGACAAGAGCAAAGTCATAAA
CTCTGCTCTGGAATTACTCAATGAAGTCGGTATCGAAGGCCTGA
CGACAAGGAAACTCGCTCAAAAGCTGGGAGTTGAGCAGCCTAC
CCTGTACTGGCACGTGAAGAACAAGCGGGCCCTGCTCGATGCC
CTGGCAATCGAGATGCTGGACAGGCATCATACCCACTTCTGCCC
CCTGGAAGGCGAGTCATGGCAAGACTTTCTGCGGAACAACGCC
AAGTCATTCCGCTGTGCTCTCCTCTCACATCGCGACGGGGCTAA
AGTGCATCTCGGCACCCGCCCAACAGAGAAACAGTACGAAACC
CTGGAAAATCAGCTCGCGTTCCTGTGTCAGCAAGGCTTCTCCCT
GGAGAACGCACTGTACGCTCTGTCCGCCGTGGGCCACTTTACAC
TGGGCTGCGTATTGGAGGATCAGGAGCATCAAGTAGCAAAAGA
GGAAAGAGAGACACCTACCACCGATTCTATGCCCCCACTTCTGA
GACAAGCAATTGAGCTGTTCGACCATCAGGGAGCCGAACCTGC
CTTCCTTTTCGGCCTGGAACTAATCATATGTGGCCTGGAGAAAC
AGCTAAAGTGCGAAAGCGGCGGGCCGGCCGACGCCCTTGACGA
TTTTGACTTAGACATGCTCCCAGCCGATGCCCTTGACGACTTTG
ACCTTGATATGCTGCCTGCTGACGCTCTTGACGATTTTGACCTTG
ACATGCTCCCCGGCTAACCCAAACCCGCTGATCCGCCCCTCTCC
CTCCCCCCCCCCTAACGTTACTGGCCGAAGCCGCTTGGAATAAG
GCCGGTGTGCGTTTGTCTATATGTTATTTTCCACCATATTGCCGT
CTTTTGGCAATGTGAGGGCCCGGAAACCTGGCCCTGTCTTCTTG
ACGAGCATTCCTAGGGGTCTTTCCCCTCTCGCCAAAGGAATGCA
AGGTCTGTTGAATGTCGTGAAGGAAGCAGTTCCTCTGGAAGCTT
CTTGAAGACAAACAACGTCTGTAGCGACCCTTTGCAGGCAGCG
GAACCCCCCACCTGGCGACAGGTGCCTCTGCGGCCAAAAGCCA
CGTGTATAAGATACACCTGCAAAGGCGGCACAACCCCAGTGCC
ACGTTGTGAGTTGGATAGTTGTGGAAAGAGTCAAATGGCTCTCC
TCAAGCGTATTCAACAAGGGGCTGAAGGATGCCCAGAAGGTAC
CCCATTGTATGGGATCTGATCTGGGGCCTCGGTGCACATGCTTT

ACATGTGTTTAGTCGAGGTTAAAAAAACGTCTAGGCCCCCCGAA
CCACGGGGACGTGGTTTTCCTTTGAAAAACACGATGATAATATG
GCCACAACCATGGGATCGGCCATTGAACAAGATGGATTGCACG
CAGGTTCTCCGGCCGCTTGGGTGGAGAGGCTATTCGGCTATGAC
TGGGCACAACAGACAATCGGCTGCTCTGATGCCGCCGTGTTCCG
GCTGTCAGCGCAGGGGCGCCCGGTTCTTTTTGTCAAGACCGACC
TGTCCGGTGCCCTGAATGAACTGCAGGACGAGGCAGCGCGGCT
ATCGTGGCTGGCCACGACGGGCGTTCCTTGCGCAGCTGTGCTCG
ACGTTGTCACTGAAGCGGGAAGGGACTGGCTGCTATTGGGCGA
AGTGCCGGGGCAGGATCTCCTGTCATCTCACCTTGCTCCTGCCG
AGAAAGTATCCATCATGGCTGATGCAATGCGGCGGCTGCATAC
GCTTGATCCGGCTACCTGCCCATTCGACCACCAAGCGAAACATC
GCATCGAGCGAGCACGTACTCGGATGGAAGCCGGTCTTGTCGA
TCAGGATGATCTGGACGAAGAGCATCAGGGGCTCGCGCCAGCC
GAACTGTTCGCCAGGCTCAAGGCGCGCATGCCCGACGGCGAGG
ATCTCGTCGTGACCCATGGCGATGCCTGCTTGCCGAATATCATG
GTGGAAAATGGCCGCTTTTCTGGATTCATCGACTGTGGCCGGCT
GGGTGTGGCGGACCGCTATCAGGACATAGCGTTGGCTACCCGT
GATATTGCTGAAGAGCTTGGCGGCGAATGGGCTGACCGCTTCCT
CGTGCTTTACGGTATCGCCGCTCCCGATTCGCAGCGCATCGCCT
TCTATCGCCTTCTTGACGAGTTCTTCTGATGATCAGCCTCGACTG
TGCCTTCTAGTTGCCAGCCATCTGTTGTTTGCCCCTCCCCCGTGC
CTTCCTTGACCCTGGAAGGTGCCACTCCCACTGTCCTTTCCTAAT
AAAATGAGGAAATTGCATCGCATTGTCTGAGTAGGTGTCATTC

**Supplemental Table 2: ColabFold settings for the generation of rHuLub C- and N-terminal domain structure.**

| Parameter | Value |
|---|---|
| num_relax | 1 |
| template_mode | pdb100 |
| msa_mode | mmseqs2_uniref_env |
| pair_mode | unpaired_paired |
| model_type | auto |
| num_recycles | auto |
| recycle_early_stop_tolerance | auto |
| pairing_strategy | greedy |
| max_MSA | auto |
| Num_seeds | 1 |

**Supplemental Table 3: APBS-PDB2PQR settings for the generation of electrostatic potential renderings.**

| Parameter | Value |
|---|---|
| Forcefield | PARSE |
| Center grid on molecule (course) | 1 |
| Center grid on molecule (fine) | 1 |
| Type of PBE to be solved | Linearized |
| Boundary condition definition | Single Debye-Huckel |
| Biomolecular dielectric constant | 2 |
| Dielectric constant of solvent | 78.54 |
| Mapping method | Cubic B-spline discretization |
| Number of grid points per Å$^2$ | 10 |
| Model for dielectric ion-accessibility coefficient | 9-point harmonic averaging |

| | |
|---|---|
| Radius of the solvent molecules | 1.4 |
| Size of support for spline-base surface definition | 0.3 |
| Temperature | 298.15 |
